# Proteome-scale amino-acid resolution footprinting of protein-binding sites in the intrinsically disordered regions of the human proteome

**DOI:** 10.1101/2021.04.13.439572

**Authors:** Caroline Benz, Muhammad Ali, Izabella Krystkowiak, Leandro Simonetti, Ahmed Sayadi, Filip Mihalic, Johanna Kliche, Eva Andersson, Per Jemth, Norman E. Davey, Ylva Ivarsson

**Author notes:** Equal contribution (Caroline Benz, Muhammad Ali, Izabella Krystkowiak).

## Abstract

Specific protein-protein interactions are central to all processes that underlie cell physiology. Numerous studies using a wide range of experimental approaches have identified tens of thousands of human protein-protein interactions. However, many interactions remain to be discovered, and low affinity, conditional and cell type-specific interactions are likely to be disproportionately under-represented. Moreover, for most known protein-protein interactions the binding regions remain uncharacterized. We previously developed proteomic peptide phage display (ProP-PD), a method for simultaneous proteome-scale identification of short linear motif (SLiM)-mediated interactions and footprinting of the binding region with amino acid resolution. Here, we describe the second-generation human disorderome (HD2), an optimized ProP-PD library that tiles all disordered regions of the human proteome and allows the screening of ~1,000,000 overlapping peptides in a single binding assay. We define guidelines for how to process, filter and rank the results and provide PepTools, a toolkit for annotation and analysis of identified hits. We uncovered 2,161 interaction pairs for 35 known SLiM-binding domains and confirmed a subset of 38 interactions by biophysical or cell-based assays. Finally, we show how the amino acid resolution binding site information can be used to pinpoint functionally important disease mutations and phosphorylation events in intrinsically disordered regions of the human proteome. The HD2 ProP-PD library paired with PepTools represents a powerful pipeline for unbiased proteome-wide discovery of SLiM-based interactions.

## Introduction

System-wide insights into protein-protein interactions (PPIs) are crucial for a comprehensive description of cellular function and organization, and a molecular understanding of genotype-to-phenotype relationships. Impressive advances are being made towards illuminating the human interactome. For example, Luck *et al*. recently provided the human reference interactome (HuRI), a map of about 53,000 human PPIs generated by yeast-two-hybrid (Y2H) all-by-all testing ^1^. Moreover, Huttlin *et al*. released BioPlex 3.0 (biophysical interactions of ORFeome-based complexes), a dataset generated through affinity-purification coupled to mass spectrometry (AP-MS) that contains close to 120,000 interactions involving more than 14,500 proteins ^2^. Nevertheless, there are some important shortcomings with these interactomics data: BioPlex ^2,3^ does not necessarily report on binary interactions but rather on larger complexes and has a bias for stable interactions, and neither of the studies provide information on the interacting regions of the protein pairs.

Here, we focus on proteome-wide screening of short linear motif (SLiM)-based interactions involving a folded domain in one protein and a short peptide present in an intrinsically disordered region (IDR) in another protein. On average, a SLiM interface buries only 3-4 residues in the binding pocket of the folded binding partner ^4^ and consequently, SLiM-based interactions are often of low-to-mid micromolar affinities ^5^. Given these unique properties, SLiM-based interactions are prevalent and crucial for dynamic processes such as cell signaling and regulation. For example, SLiMs commonly direct the transient complex association, scaffolding, modification state, half-life and localization of a protein ^6^. The Eukaryotic Linear Motif (ELM) database, which is the most comprehensive, manually curated database of SLiMs, currently holds only 2,092 experimentally validated human SLiM instances ^6^. Most of these interactions have been characterized through low-throughput experiments, as the properties that make SLiM-based interactions suited for their physiological function (transient interactions, micromolar affinities and cell state-dependent conditionality) make them difficult to capture experimentally by classical high-throughput PPI discovery methods ^7^. Many SLiM-based PPIs rely on additional binding sites present in the interacting proteins, which further complicates their identification ^8,9^. Consequently, it is likely that the majority of SLiMs remain to be discovered ^10^.

Proteomic peptide-phage display (ProP-PD) offers a high-throughput approach to simultaneously identify novel SLiM-based PPIs and the binding motifs in the IDRs at amino acid resolution (**Fig. 1**) ^11,12^. The approach can also provide the specificity determinants of the SLiM-binding bait through the analysis of enriched motif consensuses in the selected peptides. In ProP-PD, a phage-encoded peptide library is computationally designed to display the disordered regions of a target proteome as highly overlapping peptides ^11^. The ProP-PD library is used in selections against immobilized bait proteins, and the peptide-coding regions of binding enriched phage pools are analyzed by next-generation sequencing (NGS). We have previously constructed a first-generation human disorderome (HD1) ^11^, and used this resource to identify interactors and binding sites for several proteins, including the docking interactions of the phosphatases PP2A ^13^, PP4 ^14^ and calcineurin ^15^. However, the HD1 library suffers from limitations that have hampered the exploitation of the full power of the approach. These limitations stem from quality issues of the commercially obtained oligonucleotide library used to make the first-generation library, as well as from library design parameters. The field has also been limited by a lack of general guidelines on how to design ProP-PD experiments, post-process the results and attribute confidence to the selected peptides.

**Figure 1.**
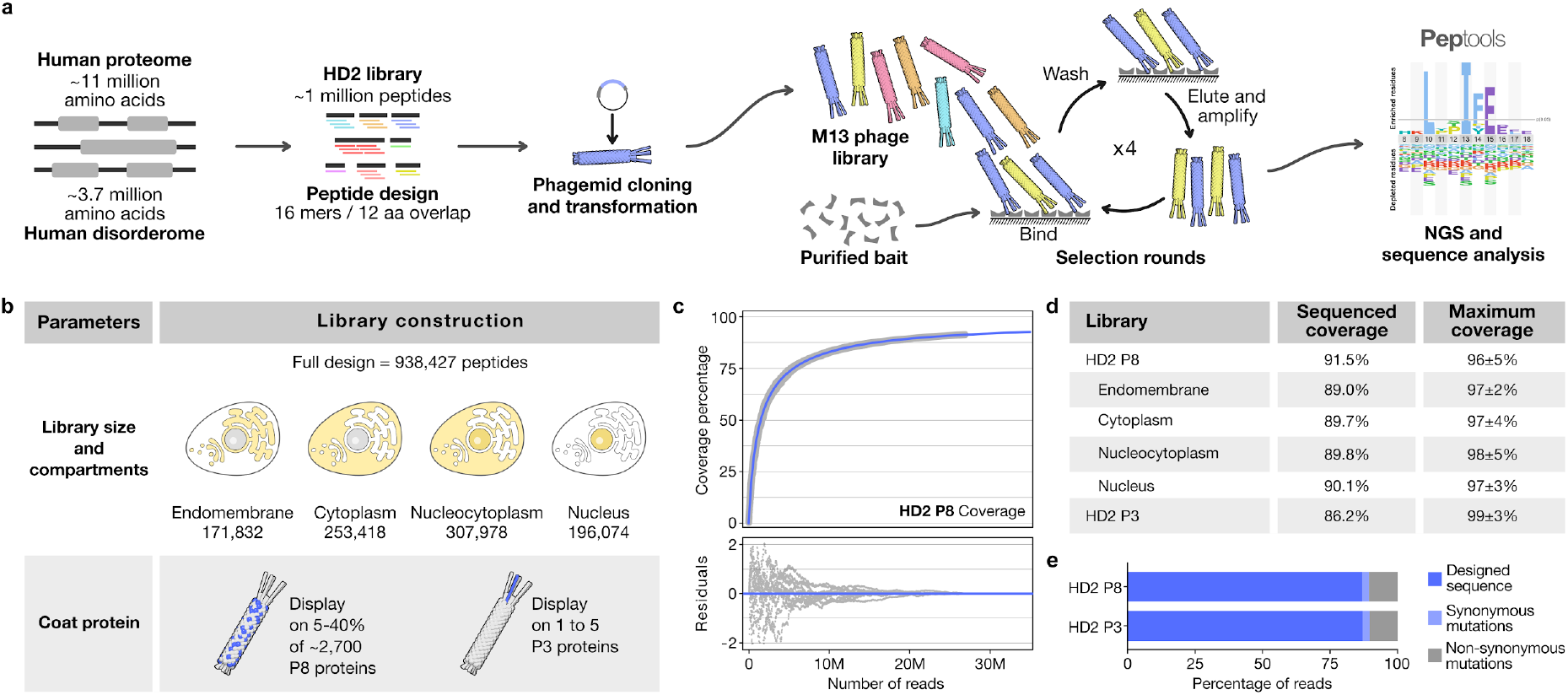
ProP-PD workflow, library design and quality. (**a**) Schematic visualization of the HD2 phage library design, cloning process, ProP-PD selection and data analysis. (**b**) Two main library parameters were explored in this work: (*i*) comparing selection results from the whole human disorderome library versus sub-libraries grouped by subcellular localization, and (*ii*) the display of the HD2 peptide library design on phage proteins P8 (multivalent, HD2 P8) and P3 (monovalent, HD2 P3), respectively. (**c**) Coverage of the HD2 P8 library as a function of sequencing depth, fitting of data to a double hyperbolic model (in blue) and residuals are shown. The contribution of reads of all sub-libraries were considered for calculating the coverage of the HD2 P8 library, as it is the result of the balanced combination of all compartment-specific sub-libraries. (**d**) Summary of observed sequence coverages and estimated maximum coverages of the constructed variant libraries (calculated as depicted in panel (c) including propagated fitting errors). (**e**) Percentage of sequences in the HD2 P8 and HD2 P3 libraries of correct length that match the library design or contain mutations.

Based on our previous experience we have developed a second-generation human disorderome library (HD2; http://slim.icr.ac.uk/phage_libraries/human/proteins.html), and an online toolkit for annotation and analysis of selected peptide ligands termed PepTools (http://slim.icr.ac.uk/tools/peptools/). We have further defined general guidelines on how to analyze the results. We evaluated the HD2 ProP-PD library by screening against 34 bait proteins representing 30 distinct domain families with known motif-mediated interaction partners listed in the ELM database ^6^. We also screened against the HEAT repeat of importin subunit beta-1 (KPNB1 HEAT), which is a challenging test case due to its typically low affinity for individual peptide ligands ^16^. We benchmarked the results against a compiled ProP-PD benchmarking set of previously validated SLiMs. Selections against the novel HD2 library captured 65 (19.3%) of the 337 known SLiM-mediated interactions for the screened protein domains. Furthermore, we uncovered 2,161 potential SLiM-mediated interactions and defined the binding sites at amino acid resolution. Biophysical characterization demonstrated that ProP-PD selections capture interactions in a broad affinity span, ranging from low nanomolar to millimolar range. Focusing on the importin subunit alpha-3 (KPNA4) binding of nuclear localization signals (NLSs), we validated the functional relevance of novel interactions through cellular localization assays. We further systematically tested phage display parameters to define the optimal analysis setup by examining the peptide selection biases of designed ProP-PD libraries and combinatorial peptide phage display libraries (*i.e*., displaying random sequences), the use of cell compartment-specific sublibraries (i.e. library designs based only on proteins localized to endomembrane, cytoplasm or nucleus), and the display on the minor coat protein P3 instead of the major coat protein P8. Finally, we explored the effects of phosphorylation or disease-related mutations on the uncovered binding sites, thus highlighting the advantage of simultaneous PPI screening and binding site identification.

## Results and discussion

### Design of the HD2 phage library

We designed a phage-encoded library of peptides representing the IDRs of the intracellular human proteome (**Fig. 1a, SI Table S1**). These disordered regions were tiled as 16 amino acid long peptides that overlap by 12 amino acids. The library contains 938,427 peptides from 16,969 proteins and covers 4,442,952 residues (~⅓ of the human proteome tiled with overlapping peptides). The library was subdivided into different, partially overlapping, pools based on the cellular localization of the peptide-containing proteins (cytoplasmic, endomembrane, cytoplasmic & nuclear, and nuclear based on localization annotation) (**Fig. 1b**) to allow for compartment specific sampling of the interaction space. Extensive information on design parameters is provided in the online methods and an interactive website to explore the full library design is available at http://slim.icr.ac.uk/phage_libraries/human/proteins.html.

### Construction and quality control of HD2 ProP-PD libraries

The designed sequences were displayed using a hybrid M13 phage system where the wild-type coat proteins are provided by the M13K07 helper phage, and the library peptides are fused to one of the coat proteins of the M13 phage. Fusion of the peptides to the P8 protein results in the display of peptides on 5-40% of the ~2,700 copies of the P8 protein on each phage ^17^. The high avidity of the P8 displayed peptides facilitates the capture of weak interactions ^18^. To evaluate how the selection results are affected by multivalent versus monovalent display we also generated a version of the HD2 library displayed on the minor coat protein P3 (HD2 P3) (**Fig. 1b**), which in the hybrid M13 phage system results in the display of one peptide per phage. We obtained oligonucleotides encoding the designed peptides flanked by phagemid annealing sites. The oligonucleotides were used to create libraries of genes coding for the designed peptides fused to the N-terminal part of the P8 or P3 protein flanked by glycine-serine linker regions. NGS of the phage libraries confirmed that ~90% of the designed peptide sequences were present in the constructed libraries, and the predicted maximum library coverage percentage surpassed 95% by non-linear regression fitting of the NGS data. (**Fig. 1 c-d**). As each amino acid of the IDRs is covered by at least two overlapping peptides, this design ensures full coverage of the human IDRs by the library. Almost 80% of the incorporated oligonucleotide sequences were of correct length and 90% of them encoded for peptides that matched the library designed (**Fig. 1e & SI Fig. S1a**), and there was no major overrepresentation of sequences in the constructed library (**SI Fig. S1b**). We thus confirmed that the constructed phage libraries have high coverage and are of high quality.

### Phage selections

The HD2 ProP-PD libraries (displayed on P8 or P3) and the first-generation HD1 library (displayed on P8) were used in triplicate selections against a glutathione transferase (GST)-tagged benchmarking set of 34 well-known SLiM-binding domains from 30 domain families (**Table 1**, **SI Table S2**). The bait proteins were chosen to cover a variety of SLiM binding pockets (**SI Fig. S2**) that recognize different types of motifs in terms of motif length, secondary structure when bound and amino acid composition (**Table 1**). In addition, we included the HEAT domain of KPNB1 that binds to FG motifs within nucleoporins with millimolar range affinities, which is ~10-100 fold weaker affinity than most of the other SLiM-based interactions tested here ^16^. GST was used as a negative control, together with a set of protein domains not expected to bind to the library peptides based on their reported binding preferences ^6^, namely the phospho-peptide binding proteins 14-3-3 protein sigma (SFN 14-3-3) ^19^ and the interferon regulatory factor 3 (IRF3 IRF-3) ^20^, the N-acetyl-peptide binding PONY domain of the DCN1-like protein 1 (DCUN1D1 PONY) ^21^, and the C-terminal binding Cap-Gly domain of CAP-Gly domaincontaining linker protein 1 (CLIP1 Cap-Gly) ^6^. As the libraries described here do not display free N-terminal or C-terminal residues, and no post-translational modifications are introduced, these domains should represent valid negative controls.

**Table 1.**
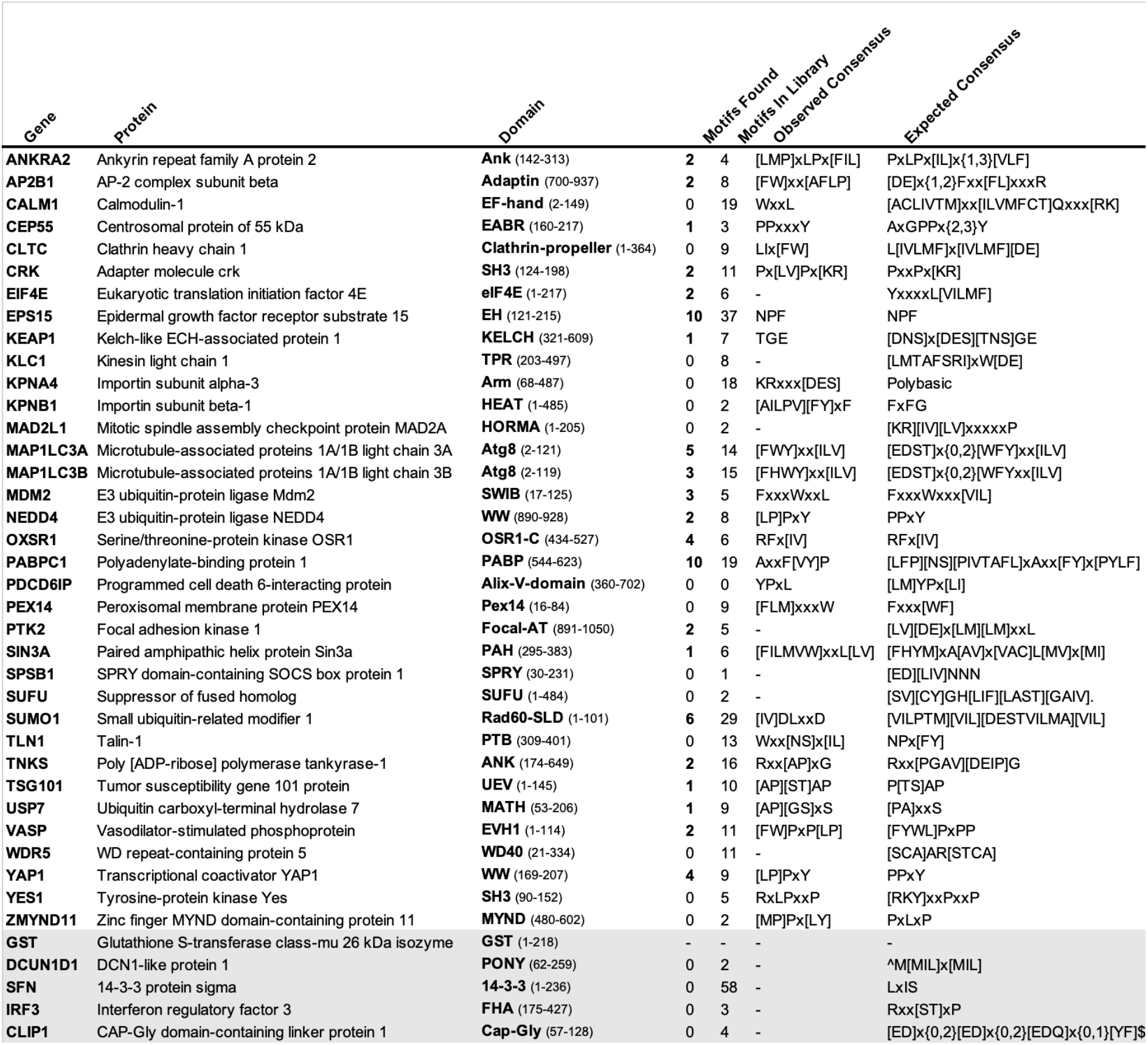
Overview of ProP-PD baits and HD2 P8 selection results. Overview of the bait constructs screen in the current study, the number of validated motifs discovered in selection for each bait, the number of validated motifs present in the HD2 library, the enriched motif consensus in the peptides selected for each bait and the expected consensus for each bait. Grey shaded area indicates baits used as negative controls.

In addition to the ProP-PD selections, we further performed selections against a combinatorial peptide phage display library with high complexity ^22^ (estimated 10^10^ diversity) to evaluate to what extent the ProP-PD library design circumvents the known issue with selection of tryptophan-rich peptides ^23^.

### Initial NGS data processing and peptide annotation

The peptide-coding regions of the resulting binding-enriched phage pools were barcoded and analyzed by NGS. The DNA sequences were translated to amino acid sequences, resulting in peptides associated with sequencing read counts. To permit comparison between screens and sequencing batches NGS counts were normalized for each peptide by the total NGS counts in each replicate selection. A merged list of peptides found through replicate screens against a given bait protein was created. For each peptide the number of replicate screens it occurred in, as well the identification of other overlapping peptides sequences were annotated (see **Fig. 2b**). The peptide sequences were mapped to the human proteome with PepTools (http://slim.icr.ac.uk/tools/peptools/), a novel web-based tool developed for the annotation of protein regions built on the annotation framework of the PSSMSearch ^24^ tool. The sets of identified replicated and/or overlapping peptides for each bait were analyzed by the SLiMFinder ^25^ consensus discovery tool for enriched specificity determinants. The consensus-containing peptides for each bait were aligned, and the alignment was converted to a position-specific scoring matrix (PSSM) using the PSI-BLAST IC method of PSSMSearch. All peptides found for a given bait were then scored by the defined PSSM, which provided a bait specificity determinant score. Finally, peptides were annotated with information related to shared annotations between the peptide containing protein and the bait protein (e.g., protein co-localization, shared functional ontology terms and previous evidence of an interaction between the bait and peptide containing protein). These data are useful for later prioritization of peptides for validation ^11,13,14^. Available information about post-translational modification (PTM) sites and disease associated single-nucleotide polymorphism (SNPs) within the peptides were also annotated. The PepTools annotated datasets generated in this project are available online (**SI Table S3**: http://slim.icr.ac.uk/data/proppd_hd2_pilot).

**Figure 2.**
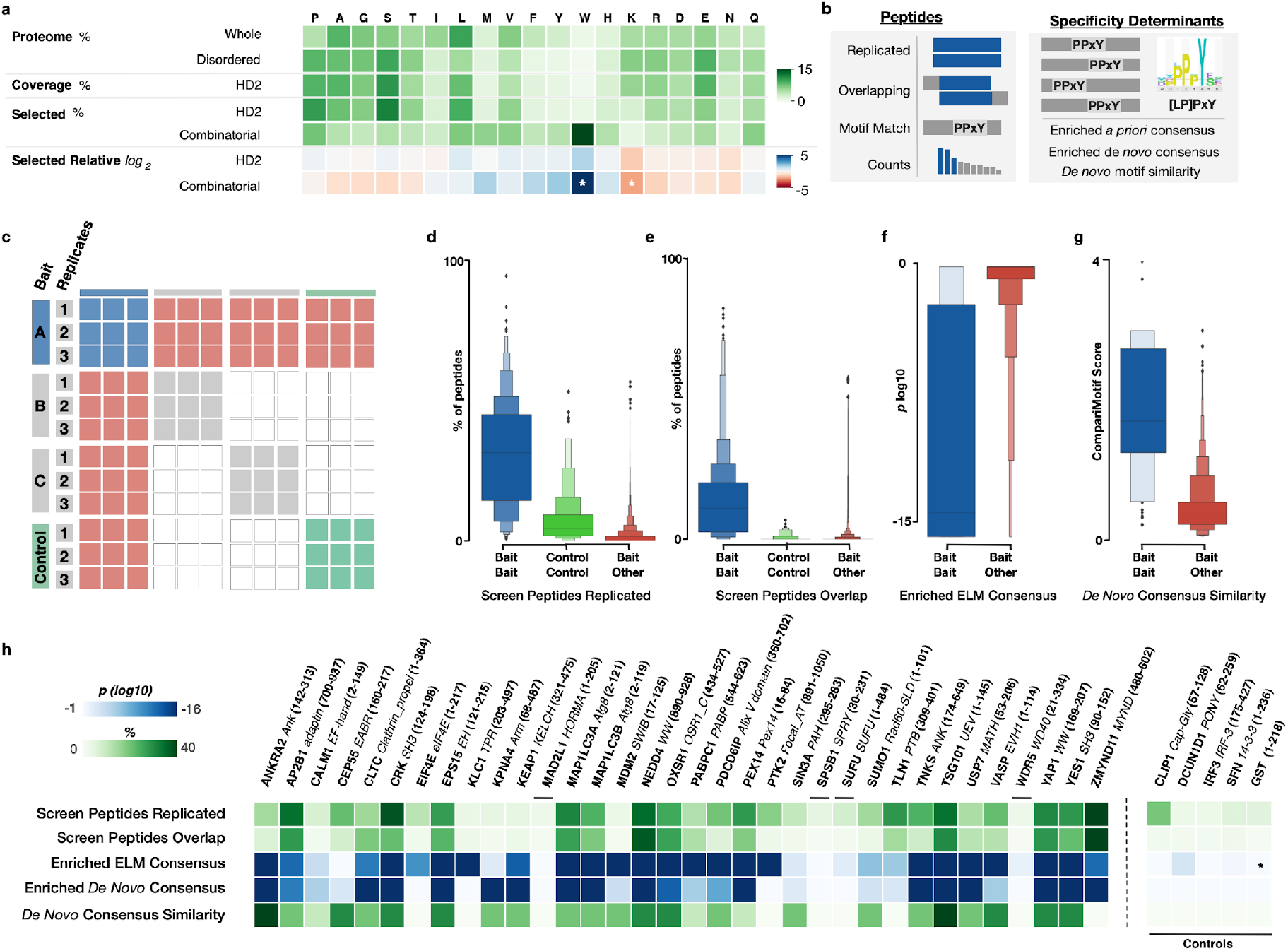
Selection quality metrics. (**a**) Amino acid frequency (green color) in *i*) the human proteome, *ii*) the predicted IDRs, *iii*) the HD2 library design, *iv*) the binding enriched phage pools from selections against the HD2 P8 library, and v) the combinatorial peptide phage display. The log2 of the relative amino acid frequencies of HD2 P8 and combinatorial peptide phage display versus the amino acid frequencies of predicted IDRs are shown in a gradient from blue to red. Note the significant enrichment of tryptophan and the depletion of lysine in the data from combinatorial peptide phage display selections (*z-score* > 2 indicated by white asterisk) but not the ProP-PD results. (**b**) Confidence metrics used in the selection quality benchmarking and peptide confidence level definitions. (**c**) Schema of the benchmarking of intra- and inter-bait replicates used in the selection quality metric benchmarking. Results of triplicate selections against a bait were compared against results from the same bait (blue), other baits (red squares), and negative controls (green squares). (**d**) Comparison of the percentage of peptides that are reproduced in pairwise comparisons between replicate selections for the same bait (blue), for the same control bait (green) and for different bait proteins (red). (**e**) Comparison of the percentage of selected peptides that are overlapping in pairwise comparisons between replicate selections for the same bait (blue), for the same control bait (green) and for different bait proteins (red). (**f**) Comparison of the log10 enrichment probability of the ELM defined motif consensus in peptides selected for the correct consensus-binding bait (blue) and all other baits (red). (**g**) Comparison of the CompariMotif similarity of the *de novo* SLiMFinder defined enriched motif in the overlapping and replicated peptides against the established ELM consensus for the bait (blue) and against all other ELM classes (red). (**h**) Selection quality metrics from panels (d) through (g) split per bait (asterisk denotes no motif defined for the bait). Enriched *de novo* consensus shows the *p value* of the SLiMFinder discovered enriched motif.

### Selection quality metrics for the HD2 P8 library

We compared the HD2 P8 data to the results of the combinatorial phage display selections. We found as previously reported ^23^ that the selection against the combinatorial phage library gave a strong bias for hydrophobic residues, and especially for tryptophan (**Fig. 2a**). The ProP-PD selection results were more similar to the proteomic frequencies of amino acids apart from minor shifts in the frequencies of tryptophan (enriched) and lysine (depleted). The HD2 library design thus largely circumvents the issue with selection of overly hydrophobic peptide sequences by combinatorial peptide phage display selection.

We evaluated the selection results globally using distinct quality metrics, namely if selected peptides were identified reproducibly in replicate screens, if we identified overlapping peptides tiling a given region, and if the peptides contained the consensus motif listed in the ELM database ^6^, or matched the *de novo* generated motifs (**Table 1, Fig. 2**). We found as expected for successful phage selections that there was an enrichment of shared peptides in datasets generated through replicate selections against the same bait proteins, but not for selections against different baits proteins (**Fig. 2c-d**). Overlapping peptides were more frequently found between data from replicate selections for the same bait as compared to unrelated screens (**Fig. 2e**). Moreover, the expected ELM consensus for a bait was often enriched in identified peptides selected for that bait (**Fig. 2f**), and the consensus motif discovered *de novo* based on the identified peptides matched the expected ELM consensus for that bait (**Fig. 2g** and **Table 1**). These observations suggest that the ProP-PD enriched peptides display the expected specificities, and that replicated peptides, overlapping peptides and enriched binding determinants are strong indicators of a successful selection. We further analyzed the results on the bait protein level (**Fig. 2h**), and found that most selections appeared successful in terms of finding binding peptides, but that the results of four of the proteins had statistics that were similar to the negative controls thus indicating that these baits failed to enrich for specific binders (**Fig. 2h**; MAD2L1, SPSB1, SUFU and WDR5).

### Evaluation of metrics for ranking of ProP-PD results and defining confidence levels

The ProP-PD derived datasets may contain enriched biophysically binding peptides, spurious peptides identified because of the depth of the sequencing and target unspecific promiscuous peptides. To enrich the data for higher confidence ligands target specific peptides should be sieved from non-relevant peptides. The four metrics that might be used to define high confidence ligands are i) reproducible occurrence in replicate selections, ii) identification of a region with overlapping peptide hits, iii) presence of a shared consensus motif, and iv) strong enrichment as indicated by high NGS read counts (**Fig. 2 and 3**). We evaluated the discriminatory power of each of the metrics using a *ProP-PD motif benchmarking dataset* (**SI Table S4;** http://slim.icr.ac.uk/data/proppd_hd2_pilot) compiled from the ELM database and structures of complexes available in the Protein Data Bank (PDB) (**Fig. 3**). The benchmarking dataset contains 337 motif instances previously reported to bind to the 34 bait proteins and that are represented in the HD2 P8 library. We found, as expected, that peptides that were discovered through the HD2 P8 selections and overlapped with the benchmarking dataset were more frequently found in replicate selections (p=2.82×10^-19^), identified with overlapping peptides (p=9.75×10^-58^) and contained the *de novo* consensus established for the ProP-PD derived peptides using SLiMFinder (p=4.41×10^-49^) (**Fig. 3b-d**). They also had higher than average normalized peptide counts (p=3.68×10^-9^). The results support that the four metrics have predictive power in terms of discriminating genuine binding peptides from the non-specific background binding events (**Fig. 3g**).

**Figure 3.**
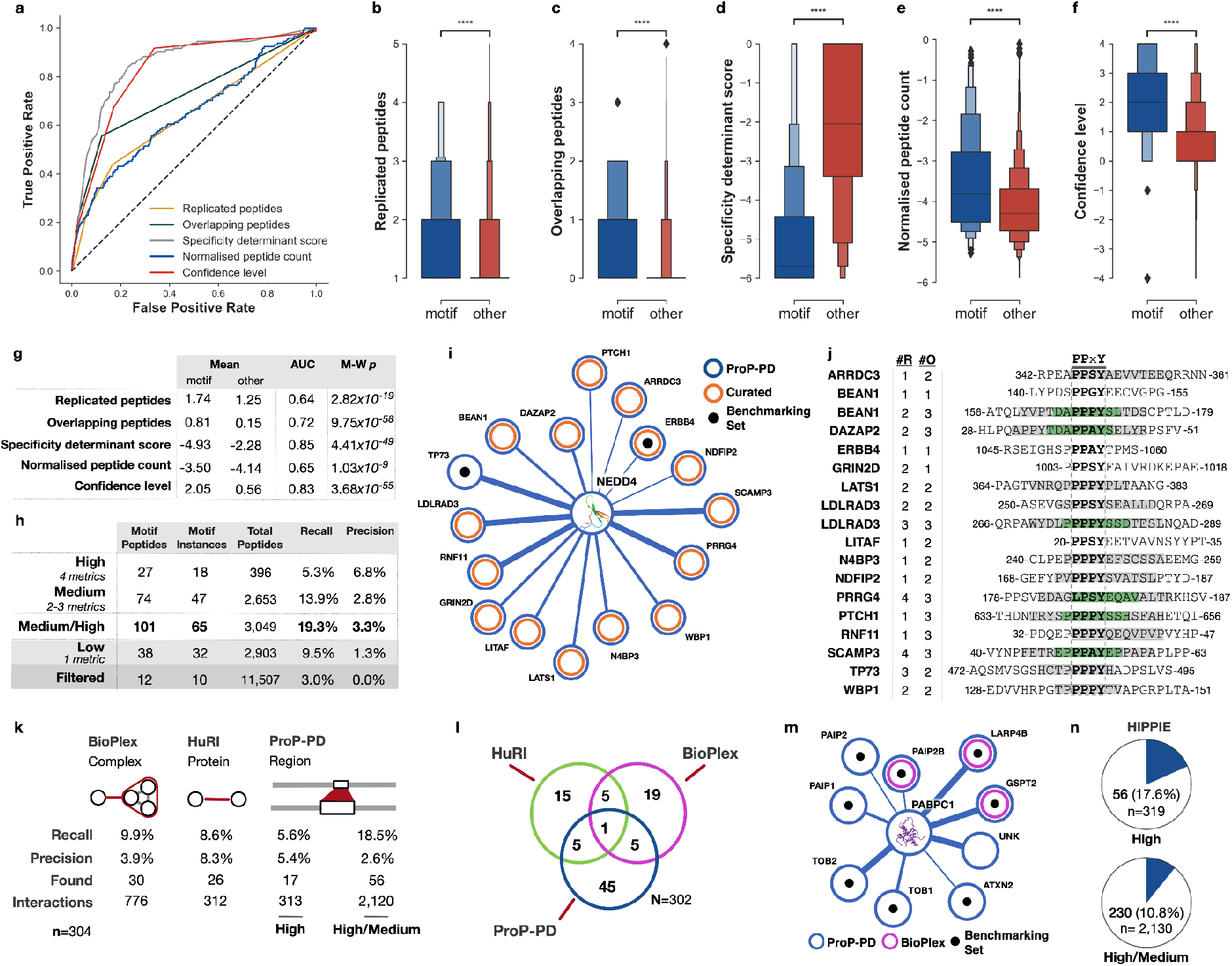
Motif rediscovery benchmarking. (**a**) ROC curves of the metrics used to assign confidence levels to peptide ligands identified through ProP-PD: i) replicated peptides, ii) overlapping peptides, iii) specificity determinant match and iv) normalized peptide count, comparing the motif-containing peptides from the ProP-PD motif benchmarking datasets to all other selected peptides (**b**) Boxplot of the number of replicated peptides for motif-containing peptides from the ProP-PD motif benchmarking datasets (blue) compared to all other selected peptides (red). (**c**) As panel b, showing overlapping peptides. (**d**) As panel b, showing the PSSM-derived specificity determinant score defining the similarity of the selected peptides to the SLiMFinder discovered enriched motif. Score is log10 of the PSSMSearch PSSM probability. (**e**) As panel b, showing log10 of the normalized peptide count. (**f**) Consensus confidence level defined based on the replicated peptides, overlapping peptides, specificity determinant match and normalized peptide count. (**g**) The predictive power, defined by the area under the ROC curve (AUC) and Mann-Whitney-Wilcoxon two-sided test with Bonferroni correction p-value (M-W p), of the four confidence metrics: replicated peptides, overlapping peptides, specificity determinant match and normalized peptide count; and the consensus confidence level metric. (**h**) Benchmarking statistics of the four consensus confidence levels and the high/medium confidence levels grouped. Recall calculated on motif instances against the benchmarking dataset of 337 motif instances. Precision calculated as the number of motif-containing peptides over the number of peptides at given confidence level. The remaining peptides are discarded. (**i**) Partial network of ProP-PD derived high/medium interactors of the fourth WW domain of NEDD4. Shown interactions involve peptides that are annotated as WW domain ligands in the ELM resource (black circles) or curated from the literature (orange circle). Line thickness indicates the number of quality metrics fulfilled by the hit (4, 3, or 2). (**j**) The ProP-PD selected peptides matching previously validated NEDD4 binding peptides from panel i annotated with the number of replicates and the overlapping peptides (grey denotes two overlapping peptides for the region and green denotes three overlapping peptides). (**k**) Interaction-centric benchmarking metrics of the ProP-PD, BioPlex and HuRI based on the 302 unique motif-mediated interactions for the 337 motif instances from the motif benchmarking dataset. *Found* is the number of motif-mediated interactions from the benchmarking dataset that were rediscovered by each method, *interactions* is the total number of interactions returned by each method for the baits in the motif benchmarking dataset. (**l**) Overlap of the validated motif interactions rediscovered by ProP-PD, BioPlex and HuRI on the interactions of the motif benchmarking dataset. (**m**) PABPC1 PPI network for proteins containing selected high and medium confidence ProP-PD peptides and annotated with BioPlex (magenta rings) and HuRI (green rings) interaction data. Edge width represents ProP-PD confidence level. Black dots represent peptides that overlap with an ELM instance known to bind the bait protein. (**n**) Overlap between the ProP-PD interactions and interactions in the HIPPIE database.

We applied two measures to benchmark the HD2 P8 results against known motif binders to our set of baits, namely *recall* (the proportion of previously validated motifs present in the library recovered through ProP-PD selections against the bait) and *precision* (the proportion of peptides found through ProP-PD selections that contain a motif previously reported to bind the bait). Cut-off values were determined for each of the four metrics through receiver operating characteristic (ROC) curve analysis (**Fig. 3a**). The resulting binary confidence criteria obtained for the individual metrics were combined for each peptide to create a single score termed ‘Confidence level’, which has four categories (‘High’, ‘Medium’, ‘Low’ and ‘Filtered’). Peptides that met all four of the metrics represent the high confidence set (396 peptides), when compared to other confidence levels these peptides have lower recall of previously validated hits but higher precision (**Fig. 3h**). Hits fulfilling two or three of the metrics (2,653) peptides) represent a medium confidence set with higher recall but lower precision. The peptides that fulfilled only one of the metrics (2,903 peptides) were considered of low confidence. Finally, a large set of peptides (11,507) that fulfilled none of the criteria contained very few previously validated ligands (10). For a stringent analysis, we focused on the 3,049 peptides in the medium/high confidence bins (**SI Table S3:** http://slim.icr.ac.uk/data/proppd_hd2_pilot). We would expect most peptides in the high confidence bin to bind the target proteins. Indeed, previous validations of HD1 P8 selection results have shown that a high proportion of peptide ligands suggested by ProP-PD are biophysical binders ^11,13–15^. In total, 65 (19.3%) of the 337 previously validated motifs in the benchmarking set were found in the medium/high confidence dataset. Performing the same analysis of the HD1 P8 data generated for the same bait proteins resulted in only 34 previously validated motif instances among 1,944 peptides, supporting that the HD2 P8 library is an improved resource for discovery of motif-based interactions.

As a final evaluation, we looked into the results of the negative controls. As expected, we identified no or few high confidence peptides in the HD2 P8 selection results for GST, IRF3 IRF-3, DCUN1D1 PONY, CLIP1 Cap-Gly or SFN 14-3-3. However, there were a few potential hits, among them the _1836-_PS**WLADIP**PWVPKDRP_-1851_ peptide from microtubule-associated protein 1A (MAP1A) that was ranked as a medium confidence ligand for SFN 14-3-3. Although less common, SFN 14-3-3 may bind to unphosphorylated ligands ^26–29^. We tested the interaction between SFN 14-3-3 and MAP1A_1836-1851_ through fluorescence-polarization (FP)-based affinity determination (**Fig. S4**), which established that it is a low affinity interaction (K_I_ 355 μM; **SI Table 5**). It can be speculated that the aspartate side chain of the MAP1A_1836-1851_ peptide mimics the negative charge of a phospho-serine, as previously shown for other 14-3-3 binding peptides ^27–29^.

### Comparison of HD2 P8 interactome to interactomics data from BioPlex and HurRI

Next, we assessed the ability of HD2 P8 phage selections to identify SLiM-based interactions by comparing the PPI data with other large-scale interactomics datasets, namely HuRI ^1^ and BioPlex ^2,3^ (**Fig. 3k-n**). As a reference set, we used the *ProP-PD motif benchmarking dataset* (**SI Table S4**) where 302 PPIs were annotated for the 337 motif instances. We used two benchmarking metrics recall and precision. In this context, recall is the proportion of PPIs from the ProP-PD motif benchmarking dataset that have been rediscovered, and precision, the proportion of re-discovered PPIs among all PPIs found. The medium/high confidence HD2 P8 data has twice the recall of BioPlex and HuRI on the motif-based interactions set, but with lower precision, particularly when compared to the HuRi data. When only the high confidence HD2 P8 data is considered, we found it to be almost comparable to the HuRI and BioPlex interaction data in terms of precision and recall, with the advantage of also providing information on the binding sites. Notably, the intersections between the three methods were low (**Fig. 3l**) suggesting that they sample different parts of the interactome, as showcased for the poly(A)-binding protein (PABP) domain of polyadenylate-binding protein 1 (PABPC1) (**Fig. 3m**). Finally, many of the interactions discovered by HD2 P8 selections have support from other studies based on the information listed in the HIPPIE database (**Fig. 3n**), which integrates and scores information on human PPIs from various sources ^30^. In conclusion, we find that HD2 ProP-PD selections generate large-scale data on motif-mediated interactions with similar quality to other high-throughput methods while also providing amino acid resolution of the binding sites.

### A closer look on the HD2 P8 data

#### High overlap between HD2 P8 data and the benchmarking datasets for PABPC1 and SUMO1

For some bait proteins, there was a remarkable overlap between the HD2 P8 selection data and previously reported binding motifs. For example, for the PABP domain of PABPC1 (**Fig. 3m**) 18 medium/high confidence peptides were identified of which all but two overlapped with previously validated motifs. This represented 10 (53%) of the 19 PABP binding peptides in the ProP-PD motif benchmarking dataset (**SI Table S4**). The remaining two peptides mapped to an overlapping 12 residue stretch found in the RING finger protein unkempt homolog (UNK; _496-_GM**NA**N**A**LP**F**Y**P**T_-506_; bold residue denote residues matching the bait consensus), which may represent a novel PABPC1 ligand. Alternatively, it may be a ligand for the homologous PABP domain from HECT E3 ubiquitin-protein ligases UBR5, which recognizes similar motifs and is functionally more closely related to UNK ^31^.

The HD2 P8 selections against SUMO1 similarly generated high quality data, resulting in 27 medium/high confidence peptides from 20 proteins (**SI Table S3**). Of these, eight peptides contained previously known SUMO interacting motifs (SIMs) from the E3 SUMO-protein ligases PIAS1/3/4 ^32^, RANBP2 ^33^, RNF4 ^34^ and RNF111 ^35^. The remaining peptides represent potential novel ligands. Among them, we noticed that the SWI/SNF-related matrix-associated actin-dependent regulator of chromatin subfamily A containing DEAD/H box 1 (SMARCAD1; _88-_GIQ**YIDL**SSDSEDVVS_-103_) has a SUMOylation site 25 amino acids upstream of the identified reverse SIM, which further supports the possibility of an additional motif mediated interface between two proteins ^36^.

#### High-quality data for WW domains but limited overlap with ProP-PD motif benchmarking dataset explained by limited literature curation

For some bait proteins, the HD2 P8 selections returned a large number of medium/high confidence hits that matched the expected consensus motif, but with no, or very limited, overlap with the ProP-PD motif benchmarking dataset (**SI Table S4**). This may in part be explained by the lack of comprehensive curation of the literature for these motif classes. For example, the IDRs of the human proteome contain 1,067 predicted [LP]PxY motifs in 900 proteins (SLiMSearch ^25^ with IUPred disorder cut off 0.4) that match the consensus recognized by class I binding WW domains. Class I binding WW domains are found in approximately 50 human proteins ^37^. We used the fourth WW domain of E3 ubiquitin-protein ligase NEDD4 (NEDD4 WW4; a widely expressed HECT type E3 ubiquitin ligase ^38^), and the first WW domain of transcriptional coactivator YAP1 (YAP1 WW1; a key regulator of the Hippo signaling ^39^) as representative cases and identified 426 medium/high confidence unique peptides that holds 435 [LP]PxY motif instances, from 245 proteins. Of these, only 5 were found in the ProP-PD benchmarking set (**SI Table S4**). To more thoroughly evaluate the quality of the results we surveyed the WW domain literature and compiled a set of 124 experimentally validated human WW domain binding PPxY motif instances (**SI Table S6**). The NEDD4 WW4 HD2 P8 selections identified 34 of the curated instances as medium/high confidence ligands, of which 19 were previously reported as NEDD4 WW domain binders (**Fig. 3i**). The selections against YAP1 WW1 identified 40 of the reported motif WW binding motifs (12 YAP1 ligands). The analysis highlights that the recall of real binders is higher than the conservative estimate provided using the ProP-PD motif benchmarking dataset.

#### Identification of an alternative calmodulin (CaM) binding motif

CaM is a ubiquitous calcium sensor that binds SLiMs upon Ca^2+^ activation. CaM binding motifs have typically high helical propensity, net positive charge and two anchor residues. They have been classified into different groups based on the distance of the anchor residues (e.g. 1-5-10 and 1-8-14) ^40,41^. Variant motifs such as the 1-4-(8/9/10) motif have also been described ^42^. CaM further binds to IQ motifs ([FILV]Qxxx[RK]Gxxx[RK]xx[FILVWY]) in its apo form ^6^. We performed selections against CaM in the presence and absence of 1 mM Ca^2+^. We expected to see an enrichment of IQ motif-containing ligands in the absence of Ca^2+^, a variety of CaM binding motifs in the presence of Ca^2+^, and a certain overlap between the sets. We found 15 medium/high confidence peptides under both conditions, and an additional set of 141 peptides only under the Ca^2+^ condition, suggesting that Ca^2+^ primed the protein for peptide binding (**SI Table S7**). The consensus motifs generated based on peptides selected under the different conditions were similar (no Ca^2+^ WxxL; 1 mM Ca^2+^ [FHW]xx[ILV]) but distinct from the classical CaM binding motifs (**Table 1**). The HD2 P8 selections thus identified CaM ligands with the less explored 1-4-(8/9/10) motif ^42^. Ligands with the longer CaM binding motifs were not captured, likely due to the minimum length of the motifs exceeding the designed peptide length of the ProP-PD library.

### Validation of SLiM-based interactions through affinity measurements

We validated a set of HD2 P8 derived interactions through FP-based affinity measurements, using a FITC-labeled probe peptide and determining the affinities (K_I_ values) for unlabeled peptides through competition. We focused on the Kelch repeats of kelch-like ECH-associated protein (KEAP1 Kelch), the E3 ubiquitin-protein ligase Mdm2 (MDM2 SWIB), the phosphotyrosine-binding (PTB) domains of talin-1 (TLN1 PTB), and the KPNB1 HEAT domain (**Fig. 4, Table 1**).

**Figure 4.**
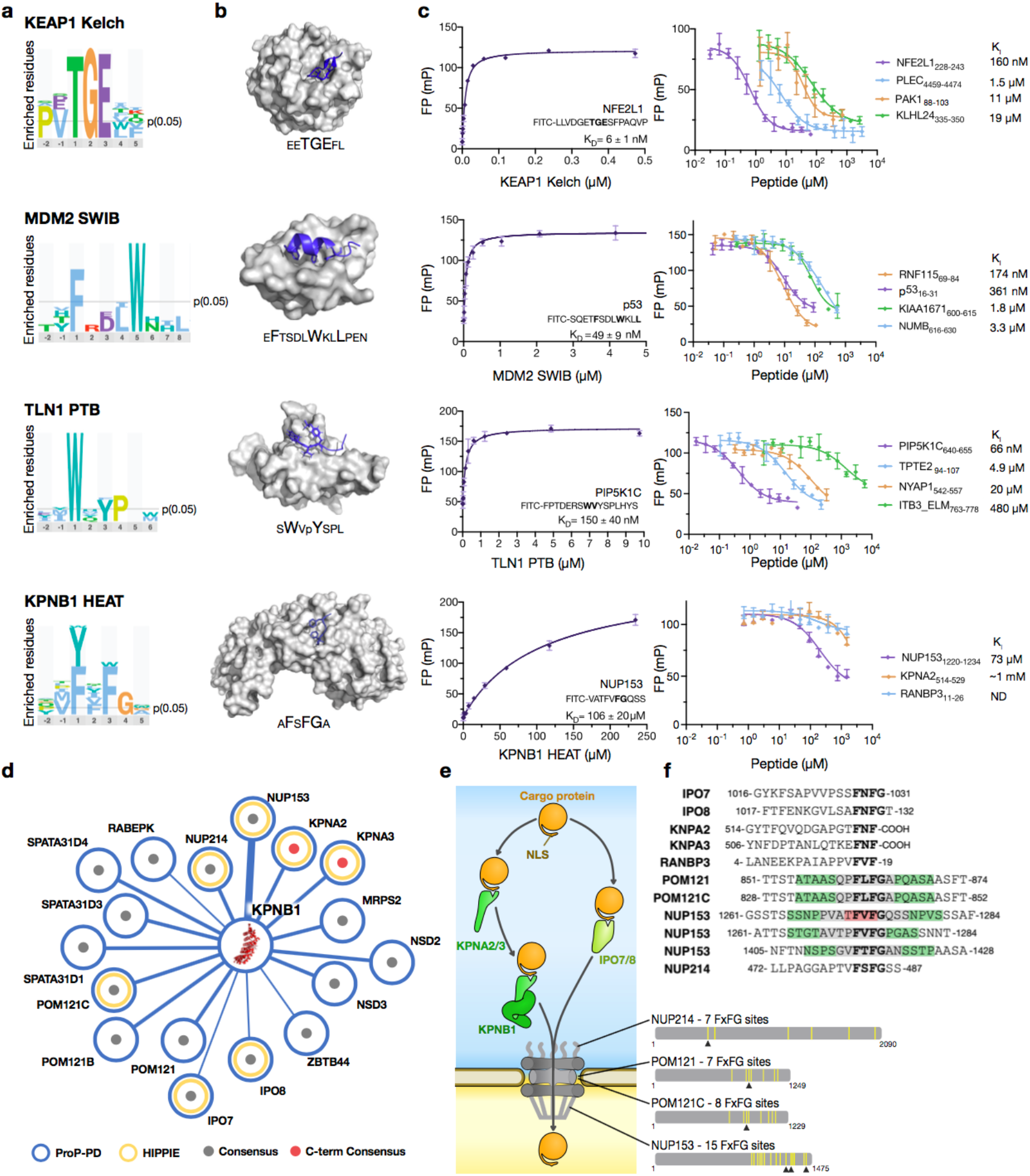
Validation of interactions by a fluorescence polarization (FP) binding assay. (**a**) Sequence logos showing the specificity determinants for the indicated bait proteins based on identified ProP-PD hits. Logos were generated using PepTools and using the medium/high confidence set of ligands as input. (**b**) Structures of KEAP1 Kelch, MDM2 SWIB, TLN1 PTB and KPNB1 HEAT with the sequences of the bound peptides indicated (PDB codes 2FLU, 1YCR, 2G35 and 1O6O). Larger amino acids indicate residues that make up the consensus motifs. **(c**) FP affinity determinations of representative hits. Affinities were measured by first determining the K_D_ value of FITC-labelled probe peptides, and then determining the affinities for unlabeled peptides through competition experiments. All experiments were performed in triplicates. See **SI Table S5** for more details. (**d**) Network of FxF(G/-coo^-^) containing KPNB1 HEAT ligands identified by HD2 selections (Edge thickness reflects the confidence level with thicker representing higher confidence level). Grey dot indicates that the peptide has a FxFG motif, red dot indicates FxF-coo^-^ motif. Annotation includes interaction data from the HIPPIE database ^28^ (yellow circle). (**e**) Schematic of KPNB1’ s role in nuclear transport together with identified FxF(G/-coo^-^) containing ligands involved in nuclear transport. The multitude of FxFG repeats in NUP213, POM121/C and NUP153 are indicated by yellow bars. Arrowheads indicate the KPNB1 binding sites identified in HD2 selections. (**f**) Sequence alignment of the KPNB1 binding peptides identified through HD2 P8 selections from proteins involved in nuclear transport (grey denotes two overlapping peptides for the region, green denotes three overlapping peptides, and red denotes four overlapping peptides).

#### KEAP1 Kelch and MDM2 SWIB bind their ligands with nano-to-micromolar affinities

KEAP1 is a substrate adaptor of the BTB-CUL3-RBX1 E3 ubiquitin ligase complex ^43^. The KEAP1 Kelch domain binds to a short negatively charged degradation motif called a degron (**Table 1, Fig. 4a**) that targets the nuclear factor erythroid 2-related factor 2 (NFE2L2) for ubiquitination and degradation. The HD2 P8 selections returned 29 medium/high confidence KEAP1 Kelch ligands from 23 proteins (**SI Table S3**) including the known ligands NFE2L1_228-243_ ^43^ and sequestosome-1 (SQSTM1_346-361_) ^44^. Affinities were determined for four peptides NFE2L1228-243, PAK1_88-103_, KLHL24_335-350_ and PLEC_4459-4474_ (**Fig. 4c, SI Table S5**), which revealed that the NFE2L1_228-243_ peptide displayed higher affinity (K_I_ 0.16 μM; **Fig. 4, Table S5**) as compared to the other peptides (1.5-19 μM K_I_ values), which may contribute to the high specificity of the protein for its primary substrate.

MDM2 mediates ubiquitination of cellular tumor suppressor p53 (p53). The SWIB domain of MDM2 binds to degrons with the consensus FxxxWxx[VIL] in target proteins ^6^. We identified 14 medium/high confidence MDM2 ligands from 12 proteins, including the known MDM2-binding peptide of p53 ^45^ (_16-_QET**F**SDL**W**KL**L**PENNV_-31_). We determined the affinities for the p53_16-31_ peptide (K_I_ of 0.36 μM), together with three novel peptide ligands from the protein numb homolog (NUMB_615-630_), the E3 ubiquitin-protein ligase RNF115 (RNF115_69-84_) and the uncharacterized protein KIAA1671 (KIAA1671_600-615_) (**Fig. 4c**). NUMB is a known substrate of MDM2 ^46^, although the interaction site has not been experimentally validated previously ^46–48^. RNF115 ubiquitinates p53 in lung adenocarcinoma ^49^, supporting a functional interplay between the two proteins. KIAA1671 is in contrast a poorly studied and largely unstructured protein, and as such belongs to what has been called as the “dark proteome”^50^. The affinities of MDM2 for the newly discovered ligands ranged from 0.17 μM for the RNF1 15_69-84_ to 3.3 μM for the NUMB_616-631_ peptide, and they are thus in similar affinity range as the known binder p53.

#### TLN1 PTB-like domain binds to ligands with a tryptophan-containing motif with high affinity

TLN1 links integrins to the actin cytoskeleton, and interacts with integrins through an interaction between the PTB-like domain of TLN1 and an NPxY motif found in the cytoplasmic tails of integrins^51^. We found 28 medium/high confidence TLN1 binding peptides from 20 proteins (**SI Table S3**), of which only one peptide contained the expected NPxY motif. Instead, the dataset was enriched in ligands with a tryptophan-containing motif (**Fig. 4**). We selected three ligands for affinity measurements: the _542-_VGPLTPL**W**TY**P**ATAAG_-557_ peptide from the neuronal tyrosine-phosphorylated phosphoinositide-3-kinase adapter 1 (NYAP1_542-557_; found in the HD1 P8 selection discussed later; **SI Table S7**), the _640-_FPTDERS**W**VY**S**PLHYS_-655_ peptide from the phosphatidylinositol 4-phosphate 5-kinase type-1 gamma (PIP5K1C_640-655_) and the _94-_DLIFTDSKLYIPLE_-107_ peptide from the phosphatidylinositol 3,4,5-trisphosphate 3-phosphatase TPTE2 (TPTE2_94-107_). We added the _763-_AKWDTAN**NP**L**Y**KEATS_-778_ peptide from integrin beta-3 (ITB3_763-778_), which is listed as a known TLN1 PTB ligand but not found among our results ^6,52^. Notably, the interaction between PIP5K1C_647-652_ and TLN1 PTB is important for targeting PIP5K1C to focal adhesions ^53^ and the structural basis for mouse TLN1 PTB and PIP5K1C has been elucidated by NMR ^54^ (PDB:2G35, **Fig. 4b**). The HD2 P8 selection thus rediscovered yet another known biologically relevant binder. While the phage derived peptides bound with affinities in the nano to micromolar range (**Fig. 4, SI Table S5**), the ITB3763-778 peptide bound weakly (K_I_ of 0.5 mM), which explains why it was not found through the phage selections.

#### Validation of KPNB1 ligands

KPNB1 plays a central role in nuclear protein import, acting both directly as a nuclear transport receptor and indirectly by associating with adapter proteins such as importin-alpha subunits (e.g., KPNA4). The KPNB1 HEAT repeat engages in low (mid-μM to mM) affinity interactions with FG repeats in proteins of the nucleoporin family ^16,55,56^. The HD2 P8 selections against KPNB1 successfully captured 23 [FWY]x[FW]G containing unique peptides found in 15 proteins (**Fig. 4d; SI Table S7**), of which 46% are found in known KPNB1 interactors that are part of the nuclear import machinery (**Fig. 4e-f**). In addition, there were two peptides with FxF-coo-motifs. We selected three of the peptides for affinity determinations, the FxFG containing nuclear pore NUP153_1120-1134_, the C-terminal motif of KPNA2_514-529_, and the RANBP_34-19_ peptide. We found that NUP153_1120-1134_ bound with a relatively high affinity for a KPNB1 interaction (70 μM K_I_) and that the C-terminal peptide of KPNA2 bound with low affinity (1 mM) (**SI Table S5**) demonstrating the large span of affinities that is recognized by KPNB1 and that can be captured through P8 HD2 selections.

#### Comparison between NGS counts and affinities

The affinity determinations demonstrated that the ProP-PD selections captured interactions with affinities within a broad affinity span. We plotted the measured affinities against the percentage of peptide counts from the NGS data to explore if there was a clear correlation between the enrichment observed and the affinities but there was no trend (**SI Fig. S5**). Hence, low count peptides may equally well represent valid targets for validations as higher count ligands. Factors such as minor biases in library composition and PCR amplification may contribute to confound affinity ranking based on NGS counts. Consequently, the method returns data that is qualitative, discriminating binders from nonbinders by enriching genuine biophysical binders from a library of almost a million peptides, however, it is not quantitative as it cannot discriminate between the small difference in affinity between these binders.

### Comparison with variant ProP-PD libraries

In addition to the HD2 P8 library we generated a variant HD2 where the designed peptides were displayed monovalently on P3, as well as HD2 P8 sub-libraries (**Fig. 1**). These libraries were used in selections against the same set of bait proteins to evaluate if monovalent presentation of the peptides or restricting the search space would result in higher quality data as measured by precision and recall. As detailed below, this was generally not the case (**Fig. 5**).

**Figure 5.**
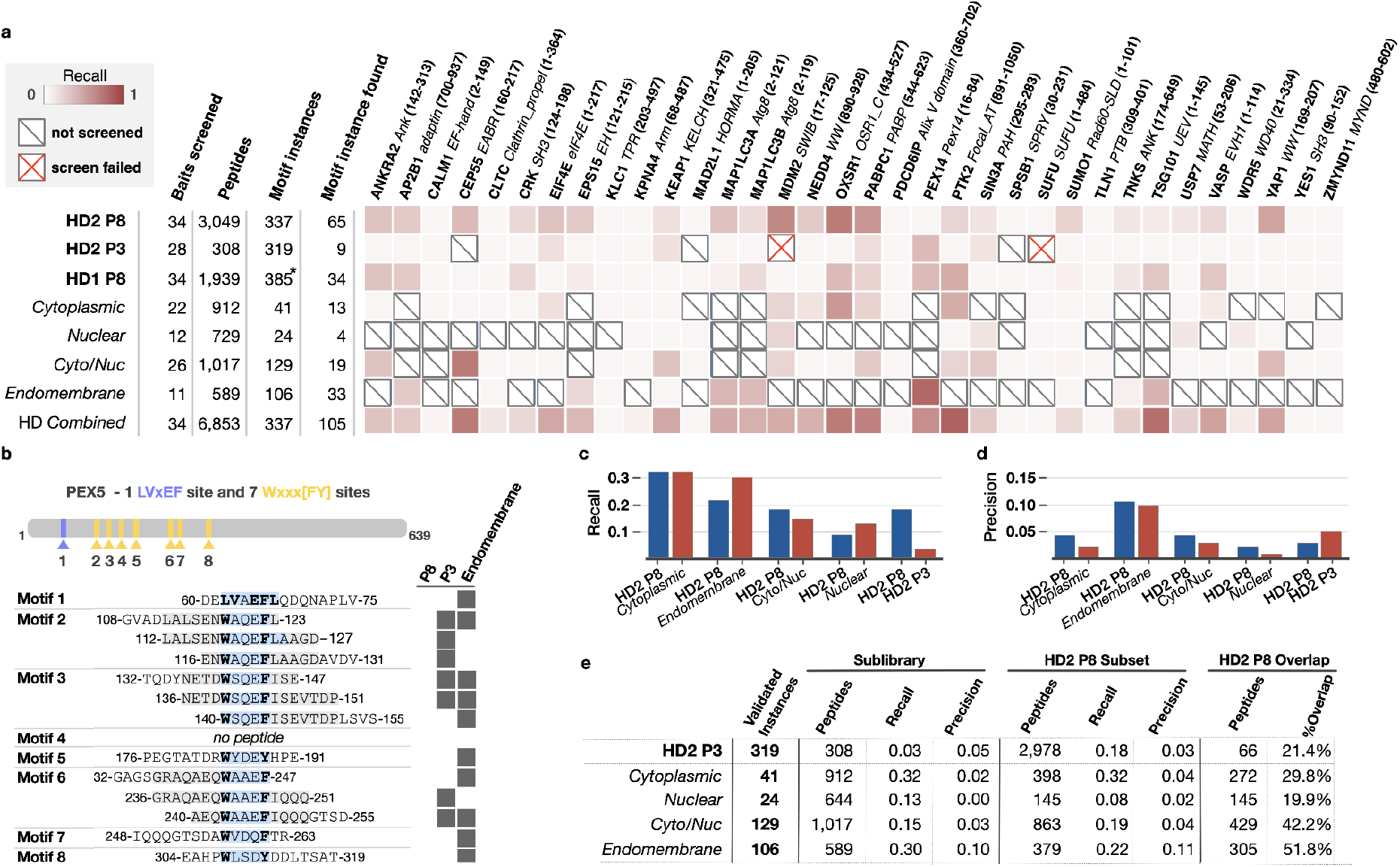
Comparison of the HD2 P8 library, the HD2 P3 library and the HD2 P8 sub-libraries. (**a**) Per bait comparison of the proportion of findable motifs in the ProP-PD motif benchmarking dataset found by each library. Overview of the pairwise comparison between the recall of the HD2 P8 library with the HD2 P8 sub-libraries and HD2 P3 library. (**b**) Overview of ProP-PD PEX14-binding peptides in PEX5 returned from different libraries (The core motif is highlighted in light blue. Motif conserved residues are in bold text). (**c**) Summary statistics of the data in panel (a) comparing the recall of the HD2 P8 library and sub-libraries or HD2 P3 library. HD2 P8 recall in the panel is calculated on the subset of motif instances that are present in the compared library. (**d**) As panel (c), summary statistics of the data in panel (a) comparing the precision of the HD2 P8 library and sublibraries or HD2 P3 library. The HD2 P8 precision in the panel is calculated on the peptides shared between the libraries being compared. (**e**) Raw data from panel (c) and (d), and overlap of the high-to-medium confidence peptides returned from the HD2 P3 library and the HD2 P8 sub-library selections with the high-to-medium confidence peptides in the HD2 P8 library screens.

#### Lower recall for HD2 P3 selection data

The HD2 P3 selections were generally less successful than the HD2 P8 selections. For two of the bait proteins (MDM2 and SUFU), selections failed to enrich for binding phage based on phage pool enzyme-linked immunosorbent assay (ELISA), suggesting that the higher avidity of the P8 display was needed to capture interactions for these bait proteins. Even when the HD2 P3 selections worked, the recall of ligands from the ProP-PD benchmarking set was generally lower (**Fig. 5a**). However, there were exceptions to the rules, such as for the N-terminal domain of the peroxisomal membrane protein PEX14 for which the HD2 P8 selections failed to return known binders, but the HD2 P3 selections returned 3 out of 8 known motif instances (**Fig. 5b; Table S7**). The HD2 P3 selection data can thus be used to reinforce and complement the HD2 P8 selection results.

#### HD2 P8 selections captured generally as many true positives as the sub-libraries

We further explored whether the use of smaller compartment specific libraries (e.g., endomembrane, nucleus, cytoplasm (**Fig. 1**) would lead to a higher recovery of previously validated hits, by avoiding competition from physiologically irrelevant ligands. However, both the recall and the precision of HD2 P8 selections benchmarked on the ProP-PD motif benchmarking dataset were on average at least as good as for the sub-library selections (**Fig. 5c-e**). There were some exceptions to the general trend, in particular related to the results from the endomembrane sub-library selections, which gave markedly better recall. A closer inspection revealed that the difference was mainly due to the results of one bait protein, namely PEX14. While the HD2 P8 selections against PEX14 returned no validated motifs, six out of the eight known PEX14-binding motifs in the peroxisomal targeting signal 1 receptor (PEX5) ^57^ were found through selection against the endomembrane sub-library (**Fig. 5b**). The PEX14 binding peptides contain a sequence pattern of acidic and hydrophobic residues (**Fig. 5c**), much like the 9 amino acids transactivation domains that interact with transcriptional regulators ^58^. When presented to the full HD2 P8 library, PEX14 is likely exposed to a large cohort of hydrophobic/acidic motifs that it would normally not see in the cell and that outcompetes the biologically relevant binders. Thus, for exceptional cases such as PEX14, screening against compartment-specific HD2 P8 sub-libraries can enhance the chances of identifying biologically relevant ligands. We conclude that for most cases there is no need to use compartment specific sub-libraries, but that they may provide an advantage for certain types of binding pockets and motif combinations, for which the inherent specificity is relatively low.

#### Reproducibility of HD2 selections

We next analyzed the high/medium confidence peptides selected for each sub-library to evaluate the reproducibility of the method (**Fig. 5e**). Of the 3,049 high/medium confidence HD2 P8 peptides identified, 1,008 (33.1%) were confirmed by selection against either a sub-library or the HD2 P3 library. For the HD2 P3 library, 22.0% (67 of 304) of the high/medium confidence peptides were confirmed by selections against one or more of the P8 libraries. In total, 1,050 peptides were reproducibly found in two or more datasets. This corresponds to 777 motif-based PPIs of which 149 have been previously observed by complementary PPI discovery methods (**SI Table S7**).

### From binding to function: validation of nuclear localization signals

To take the validation of the results from binding to function, we turned to KPNA4 that binds to NLSs and transports cargo through the NPC by the classical nuclear import pathway (**Fig. 6**). KPNA4 has two distinct NLS binding pockets (major, ARM 2-4; minor, ARM 6-8; **Fig. 6b**) ^59^. Bipartite NLSs are about 17-19 amino acids long and engage both pockets. Monopartite NLSs are typically short basic stretches, divided into five classes ^60^: class I KR[KR]R or K[KR]RK, class II [PR]xxKR[^DE] [KR] (where ^ indicates “not”); class III KRx[WFY]xxAF, class IV [RP]xxKR[KR][^DE] and class V LGKR[KR][WFY]. The class I and class II motifs overlap, and preferentially interact with the major pocket, and to a lesser extent with the minor pocket ^60^. In contrast, class III and class IV ligands have been suggested to preferentially bind to the minor pocket. Short unclassified NLSs have also been found to bind to the minor pocket, such as _273-_GSIIRKWN_-280_ from the human scramblase 4 (PLSCR4) and the _284-_KRKH_-287_ stretch from *Xenopus laevis* targeting protein for Xklp2-A (TPX2) ^61,62^.

**Figure 6.**
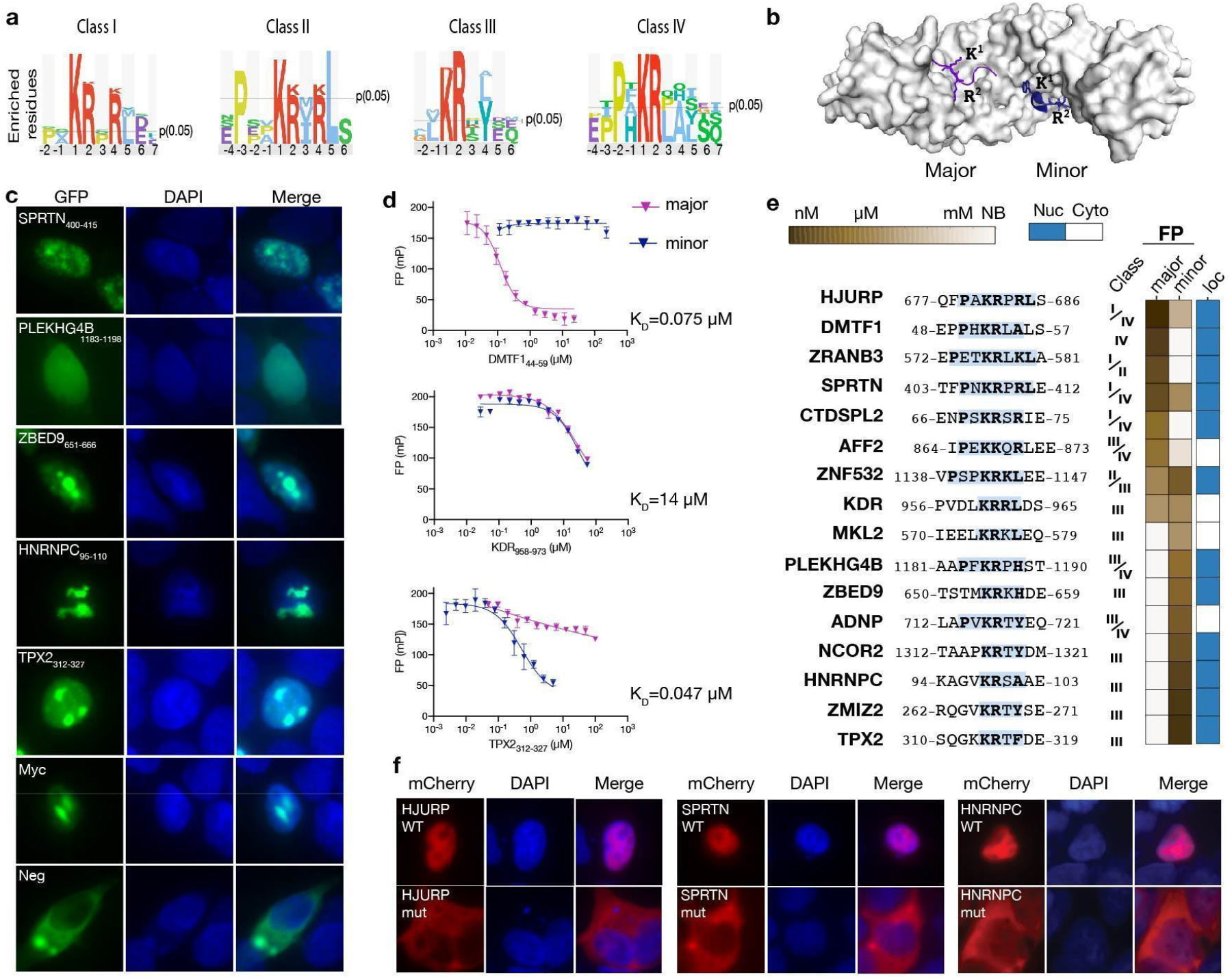
Validation of KPNA4 binding sequences as NLSs. (**a**) Sequence logos matching different NLS classes generated based on KPNA4 ProP-PD results generated using PepTools, and medium/high confidence ligands as input. (**b**) Structure of KPNA2 (PDB: 1PJN, minor groove peptide PDB:3ZIP) with ligands bound to the major (purple) and minor groove (blue). (**c**) Representative examples of the cellular localization experiment. HEK293 cells were transiently transfected with the NLS sensor and fixed 36 h after transfection, and imaged using epifluorescence microscopy. The nucleus was stained with DAPI. (**d**) FP competition experiments using FITC-Myc_320-328_ as a probe for the major groove (blue) or FITC-NCOR2_1307-1322_ as a probe for the minor groove and competing with unlabeled DMTF1_44-59_, KDR_958-973_ and TPX2_312-327_ peptides. (**e**) Core sequences of tested NLSs together with the outcome of the affinity measurement through FP and localization of the GFP-tagged peptides (see SI Fig. S7 for details). (**f**) Mutational analysis of identified NLSs in the context of full-length proteins using mCherry tagged HJURP, SPRTN and HNRNPC.

Using KPNA4 as a bait against the HD2 P8 library identified 33 peptides found in 32 proteins, of which 10 contained class I motifs (KRx[KR]) and 16 contained class III like motif (KRx[YLA]) (**SI Table S3**). To gain additional information we combined the data generated for KPNA4 using different libraries (HD2 P8, HD2 P3, HD1 P8, and the nuclear/cytoplasmic and nuclear sub-libraries) resulting in a set of 92 peptides containing NLS-like sequences found in 76 proteins. In addition to class I-IV NLS like sequences (**Fig. 6a**) four peptides contained a KRxH motif, similar to the _284-_KRKH_-287_ peptide of TPX2 (for example see **Fig. 6e**). We found prior validations supporting NLS function for 7 of the sequences ^63–67^, including the bipartite NLS of the activity-dependent neuroprotector homeobox protein (ADNP) (_711-_SLAPV**KR**T**Y**EQMEFPL_-726_ and _719-_YEQMEFPLL**KK**R**K**LDD_-734_) ^65^. We determined the affinities for 16 peptides and established their pocket specificity using FP competition experiments. We first determined the affinities of FITC-labeled probe peptides (**SI Fig. S6**), using the _320-_**P**AA**KR**V**K**LD_-328_ peptide from the Myc proto-oncogene protein (Myc) ^79^ as a probe for the major pocket, and the _1307-_P**KR**T**Y**DMMEGRVGRAI_-1322_ peptide from the nuclear receptor corepressor 2 (NCOR2) as a probe for the minor pocket. We then outcompeted the pocket specific probe peptides using unlabeled peptides (**Fig. 6d-e**, **SI Fig. S6**), and found that the 16 tested peptides bound to KPNA4 with a broad range of affinities (nM to mM) and with distinct pocket specificities. Eight peptides with class III motif only competed for the probe for the minor pocket. Similarly, the KRxH containing peptides competed only for the probe for the minor pocket, suggesting that the class III NLSs could be updated to KRx[YLAH]). Five peptides outcompete both probe peptides, which may be explained by the presence of overlapping motifs that matched both a class II and a class III motif (e.g., the _1135-_**P**SP**KR**K**L**_-1139_ stretch in ZNF532; **Fig. 6**), and to some degree by cross specificity of the two binding pockets.

We validated the function of 16 of the putative NLSs by fusing the peptides to a trimer of GFP, which gave a clear cytoplasmic localization of the negative controls, and nuclear localizations of the positive control (**Fig. 6c; SI Fig. S7**). Using this NLS sensor, we confirmed that 12 out of 16 tested peptides were functional NLSs. The two lowest affinity ligands (KDR_958-973_ and MKL2_572-587_) failed to function as NLSs, suggesting a correlation between affinity and NLS function. One of the peptides that failed to target GFP to the nucleus was the partial bipartite NLS from ADNP ^72^, which likely requires the context of the full bipartite NLS to be functional. We further tested the function of three NLSs in the context of the full-length proteins (heterogeneous nuclear ribonucleoproteins C1/C2 (HNRNPC) _95-_AGV**KR**S**A**AEMYGSVTE_-110_; DNA-dependent metalloprotease SPRTN (SPRTN) _400-_SEDTFPN**KR**P**R**LEDKTV_-416_; Holliday junction recognition protein (HJURP) _675-_DHQFPA**KR**P**R**LSEPQG-_690_). We expressed mCherry-tagged wild-type proteins and NLS mutants thereof (HNRNPC _98-_**KR/AA**_-99_; SPRTN _407-_**KR/AA**_-408_; HJURP _681-_**KR/AA**_-682_) in HEK293 cells and found that while the wild-type HJURP, HNRNPC and SPRTN proteins were efficiently targeted to the nucleus, the mutants were retained in the cytoplasm, thus supporting the importance of the identified NLSs in targeting the proteins to the nucleus (**Fig. 6f**). The ProP-PD selections thus successfully identified KNPA4 binding peptides that act as functional NLSs.

### Exploring the effects of mutations and phosphorylation on the motif-based PPI network

The medium/high confidence data from selections against the HD2 P8 library, the HD2 P8 sub-libraries and the HD2 P3 library were combined to provide an extensive network of PPIs with amino acid resolution information of the binding sites. The PPI network was annotated with a variety of biologically relevant information using the PepTools server (**Table S7**). These data provided support for the biologically relevance of many ligands based on the overlap of the contextual information of the bait and prey (shared complex, localization and functional terms).

The PPI network annotation can, for example, be used to identify binding interfaces that overlap with disease mutations or phosphosites and the information is provided as a part of the PepTools annotations. We found 183 high/medium confidence peptides with 313 unique mutations, at 253 sites (**SI Fig. S8, SI Table S8**). Among the mutational information, we find for example missense mutations of the tankyrase binding peptide from the SH3 binding protein 2 (SH3BP2 _408-_PQLPHLQ**<u>R</u>**SP**<u>P</u>D<u>G</u>**QSF. 423; affected residues underlined) that are known to abrogate the interaction with TNKs and are associated with cherubism ^68–70^. We selected the two proteins KPNA4 and KEAP1 (**Fig. 7a**) to explore the effects of disease-related mutations and tested the effects of four disease associated mutations: a conservative K194R mutation of the KPNA4 binding peptide from the homeobox protein Nkx-2.5 (NKX2-5) that is associated with atrial septal defect 7 ^71–7^, two mutations of the KEAP1 binding motif of NFE2L2 (E79K and T80K) that are linked to early onset of multisystem disorder ^75^, and an R4466C mutation in the flanking region of the KEAP1 binding motif in PLEC that is linked to epidermolysis bullosa simplex although with uncertain significance on pathogenicity ^76^. The conservative K194R mutation of NKX2-5_192-207_ resulted in a striking 10-fold loss of affinity of KPNA4 binding (**Fig. 7b**), and both NFE2L2 mutants conferred a more than 1,000-fold loss of affinity. In contrast the R4466C mutation of PLEC_4457-4471_ had no effect, as could be expected given that it occurred outside of the core motif. The analysis demonstrates how the proteome-scale amino-acid resolution footprinting of protein-binding sites in the IDRs combined with the annotations provided by PepTools can be used to pinpoint the effects of disease associated mutations (**Fig. 7c-d**).

**Figure 7.**
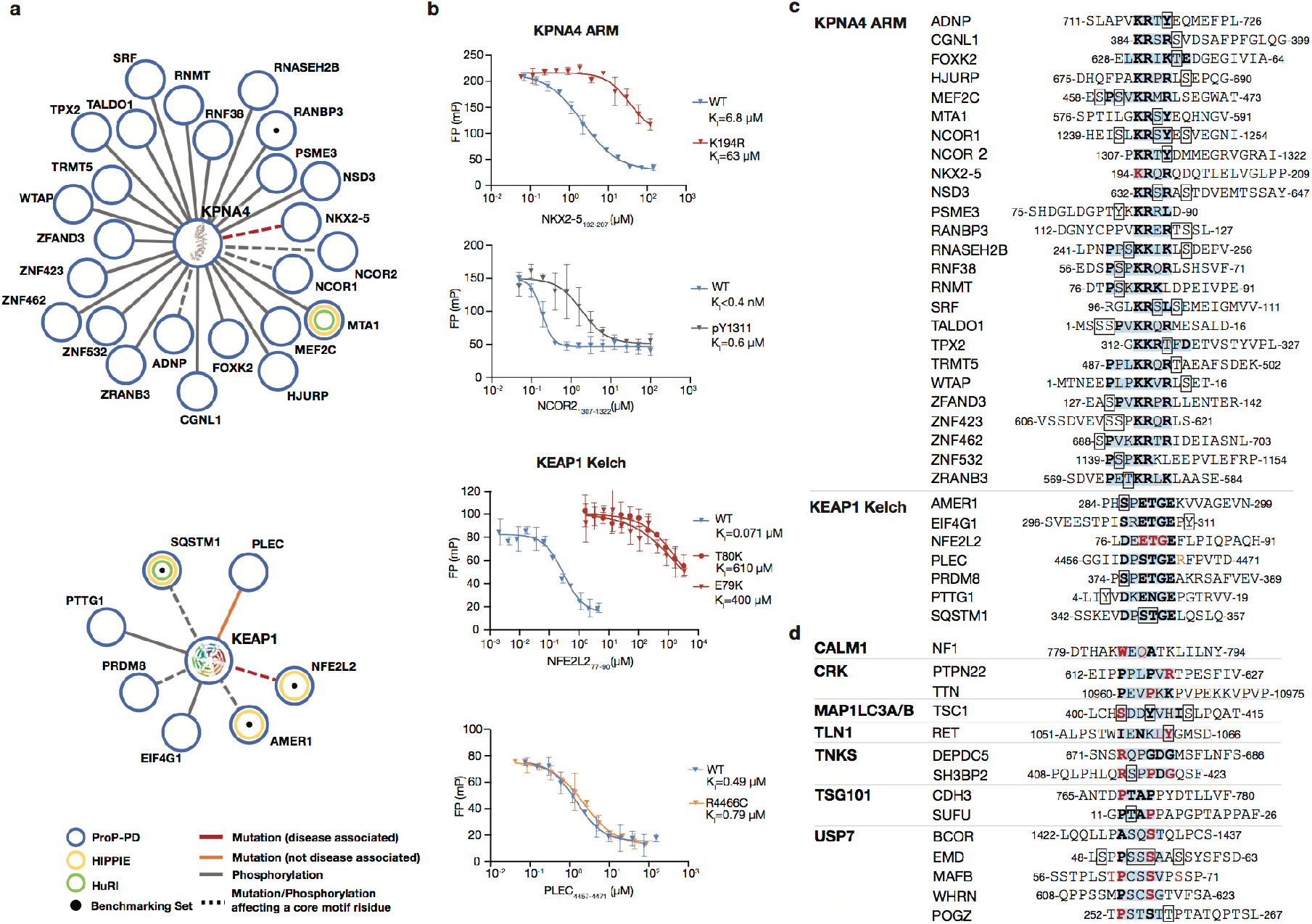
Exploring the effects of disease-related mutations and phosphorylation on uncovered interactions. (**a**) PPI networks of KPNA4 and KEAP1 showing reproducibly selected high/medium confidence interactions with disease-associated mutations, or phosphorylation sites, overlapping with the binding peptides. Only peptides with mutations or phosphorylation residues occurring in the core motif or in the flanking regions (+/- 2 residues) are shown. Mutations associated with diseases are colored in red (orange if they are not disease associated). Phosphorylated sites are colored in gray. Dashed-edge lines represent that mutations or phosphorylation sites are affecting core motif residues (**b**) FP competition experiments of wild-type, disease mutant and phospho-peptides binding to NCOR2 or KPNA4. The affinities of NXK2-5 wild-type and K194R mutant for KPNA4 were determined using a probe for major pocket (FITC-Myc_320-328_); the affinities of unphosphorylated and phosphorylated NCOR2 peptides were determined using a probe for the minor pocket (FITC-NCOR2_1307-1322_); the binding to KEAP1 were measured using FITC-NFE2L1_228-243_. (**c**) Peptide sequences related to the interactions shown in panel (a). (**d**) Additional reproducibly selected high/medium confidence peptides with disease-associated mutations in the consensus binding motif. For (c) and (d): Motif-containing regions are highlighted with blue background, motifs are indicated in bold letters, phosphorylation sites within or in vicinity of the motifs are indicated by a box, and disease associated mutations are indicated in red bold letters (mutation of motif residue).

In addition to disease mutations, we found 6,724 PTM sites in 2,755 high/medium confidence unique peptides, with phosphorylation being the most common PTM (5,868; **SI Table S9**). The results can thus also be used to identify phospho-switches that may tune binding. We experimentally tested the effect of one of these phospho-sites in the class III NLS of NCOR2 (pY1311), and found it to confer a marked loss of affinity for KPNA4, although the affinity of the NCOR2_1307-1322_ and KPNA4 is still high (Fig. 7b). Given the high affinity of the NCOR2 peptide, it can be speculated that the interaction may serve an inhibitory function, which might be relaxed by phosphorylation.

### Concluding remarks

We present a powerful experimental and bioinformatics pipeline for the proteome-scale discovery of motif-based interactions that generates results of similar quality to approaches such as AP-MS and Y2H. However, in addition ProP-PD provides amino acid resolution information on the binding sites. An added advantage is that there is no bias for interactions with highly expressed proteins, or cell type or cell cycle dependence as the selections are performed *in vitro* and bait proteins are challenged with the full IDRs of the human proteome. We acknowledge that the results are solely based on the *in vitro* interactions between isolated domains and short peptides of target proteins, and as for other large-scale interaction data, detailed validations at the level of full-length proteins and in a cellular setting are required before reaching any conclusion about the functional relevance of identified interactions. One of the fixed parameters of this study was the length of the displayed peptides, which should be sufficient to cover most of the SLiMs, given that they are typically between 3-10 amino acids ^4^ However, we note that the peptides are too short to capture certain motif classes, for example, bipartite NLSs, or the CaM IQ motifs. A future direction may thus be to create libraries expressing longer peptides, as previously done using T7 phage ^82^, which may require display on the minor coat protein P3 of the M13 phage as the display of longer peptides on the major coat protein P8 might destabilize the phage coat.

Among the challenges not addressed in this study are those related to the identification of SLiM based interactions involving PTMs, such as phosphorylation. These challenges might be addressed by, for example, the use of phosphomimetic mutations, by treating phage libraries with enzymes ^77^, or by using an expanded genetic code ^78,79^. Nevertheless, the annotations provided by PepTools can suggest potential regulation by PTMs. PepTools also provides information on disease-associated mutations in identified peptides, which can give clues about the underlying molecular determinants of diseases. Similarly, we uncovered novel interactions for known drug targets thereby improving our understanding of the therapeutically targeted proteome. Fifty of the discovered motif-containing proteins have at least one approved drug (**SI Table S7**). For two of the screened baits, MDM2 and KEAP1, small molecules have been developed to therapeutically target the motif-binding pocket ^80,81^ and the newly discovered ligands of these proteins may help explain off-target effects of these inhibitors.

In conclusion, we provide more than 2,000 human PPIs, with amino acid resolution of the binding site. We foresee that ProP-PD will contribute to the mapping of the human interactome proteome over the next decade, which will provide detailed information on binding motifs and contribute to a deeper understanding of genotype-to-phenotype relationships. What we can note from this study is that 471 of the interactions reported here are with proteins classified as poorly studied proteins in the Pharos database ^83^ and we thus shed light on understudied parts of the proteome. Given that there are more than 200 known families of SLiM-binding domains and in the range of 100,000 motif-based interactions to uncover, there is a sizable task ahead for the scientific community ^10^. Through the proteome-scale amino acid resolution footprinting offered by ProP-PD we hope to contribute insights into a considerable part of these interactions over the years to come.

## Acknowledgements

This work was supported by the Swedish Foundation for Strategic research (YI, PJ: SB16-0039), the Swedish Research Council (YI: 2016-04965; PJ:2016-04134) and Cancer Research UK (CRUK) (ND: Senior Cancer Research Fellowship C68484/A28159). Sequencing was performed by the SNP&SEQ Technology Platform in Uppsala. The facility is part of the National Genomic Infrastructure (NGI) Sweden and Science for Life Laboratory. The SNP&SEQ Platform is also supported by the Swedish Research Council and the Knut and Alice Wallenberg Foundation. We thank Prof. Ola Söderberg for access to microscope and advice related to image analysis and Dr. Evangelia Petsalaki for useful discussions related to benchmarking. We thank Lara Cardinaels, Margot Dierickx and Karl Nordström for technical support. Plasmids for expression of GST-tagged MAP1LC3A, MAP1LC3B and SUMO1 and the P3 phagemid were kindly provided by Dr. Andreas Ernst. The plasmids for expression of GST-tagged TLN1 PTB, 14-3-3 SFN, TSG101 UEV, VASP EVH1 and IRF3 IRF-3, and the combinatorial phage library were generously provided by Prof. Sachdev Sidhu. Plasmids for expression of GST-tagged NEDD4 WW4 and YAP1 WW1 were provided by Prof. Marius Sudol and Prof. Sachdev Sidhu. mCherry2-C1 was a gift from Michael Davidson. We thank Dr. Philip M Kim for his support of the project, and Dr. Peter Pryciak, Dr. Lucia Chemes and Prof. Martha Cyert for their valuable feedback on the manuscript.

## Author contribution

CB, MA, FM, EA and JK purified proteins. CB and MA constructed phage libraries. CB and JK performed phage selections. CB, FM and JK performed affinity measurements and analyzed the data. MA performed cell-based validations. LS built an NGS data analysis pipeline. IK constructed the PepTool website. ND and LS processed ProP-PD data with support of IK. ND and AS benchmarked the ProP-PD data. ND designed the custom oligonucleotide libraries with support of LS. CB, MA, JK, FM, PJ, ND and YI designed experiments. ND & YI conceived the study. CB, AM, IK, LS, AS, PJ, ND & YI wrote the paper with input from all authors.

## Data availability

The ProP-PD data is available in the supplementary materials and online at http://slim.icr.ac.uk/data/proppd_hd2_pilot. The PPI data will be deposited to the IMEX consortium through IntAct. An interactive website to explore the full library design is available at http://slim.icr.ac.uk/phage_libraries/human/. The PepTools analysis tool is available at http://slim.icr.ac.uk/tools/peptools/. All scripts used for the demultiplexing and analysis of the sequences are available at https://bitbucket.org/daveylab/phage_display_pipeline/.

## Supporting information (SI)

### SI Figures

**SI Figure S1. Non-design components and count distribution of the libraries.** (**a**) Percentages of reads obtained by NGS of the libraries generated in this work associated with sequences that matched the designed sequences, those that presented mutations either silent or non-silent, and those that due to frame-shifting, insertion or deletion mutations did not match the expected oligonucleotide length. (**b**) Distribution of NGS counts for the designed oligonucleotides in the HD2 P8 and HD2 P3 libraries.

**SI Figure S2.** Representative structures of the bait proteins domains with bound ligands and indicated binding motifs.

**SI Figure S3. NGS reads for binding enriched phage pools.** Distribution of the NGS reads that matched the peptides encoded by the library are shown grouped by bait. Control baits are shown in gray.

**SI Figure S4.** FP affinity measurements of the interactions between 14-3-3 SFN and (a) the probe peptide phospho-RAF1_255-264_ (FITC-QRSTpSTPNVH) and MAP1A_1836-1851_.

**SI Figure S5: Analysis of correlation between affinity and ProP-OD results:** Affinities were correlated to IC_50_, K_I_ and overlaps of the peptides in selections and the occurrence of peptides in selection in the different duplicates. A KPNA4 outlier was removed for better visualization.

**SI Figure S6. FP affinity measurements of KPNA4.** (**a**) Saturation curves with probes for the for the major groove (Myc_320-328_ FITC-PAAKRVKLD; purple) and minor groove (NCOR2_1307-1322_; FITC-PKRTYDMMEGRVGRAI (blue)). (**b**) FP competition binding experiment of 18 peptides against the major groove probe FITC-Myc_320-328_ (purple) and the minor groove FITC-NCOR2_1307-1322_ (blue). Unlabeled peptides used for competition: HJURP_675-690_, DMTF1_44-59_, ZRANB3_571-586_, SPRTN_400-416_, CTDSPL_64-79_, AFF2_864-879_, ZNF532_1139-1154_, KDR_958-973_, MLK2_572-587_, PLEKH64B_1183-1198_ ZBED9_651-666_, ADNP_711-726_, NCOR2_1307-1322_, HNRPNC_95-110_, ZMIZ2_264-279_, TPX2_312-327_.

**SI Figure S7: Validation of KPNA4 binding sequences as NLSs.** (**a**) Schematic of the construct used as NLS sensor. (**b**) Cellular localization of NLS sensor with different peptides inserted. HEK293 cells were transiently transfected with the NLS sensor and fixed 36 h after transfection, and imaged using epifluorescence microscopy. The nucleus was stained with DAPI. (**c**) Overview of the peptide sequences fused to the NLS sensor, with the resulting cellular localization indicated.

**SI Figure S8. Additional PPI networks.** (**a**) Networks showing interactions of reproducibly selected high/medium confidence peptides with overlapping disease-associated mutations or phosphorylation. Peptides with mutations or phosphorylation residues occurring in the core motif or in the flanking regions (+/- 2 residues) are shown. Mutations associated with diseases are colored in red (orange if they are not disease associated). Phosphorylated sites are colored in gray. Dashed-edge lines represent core motif residues holding a mutation or a phosphorylation site. (**b**) Sequences of high/medium confidence peptides with disease-associated mutations in the binding motif. Motif-containing regions are highlighted with blue background, core motifs are indicated in bold letters, phosphorylation sites within or in vicinity of the motifs are indicated by a box, and disease associated mutations are indicated in red bold letters (mutation of core residue).

### SI Tables

**SI Table S1. Library design statistics.**

Overview of the general parameters for the libraries generated for this work. For each library the number of designed oligonucleotides and their resulting number of unique peptides after translation is shown, together with redundancy percentages (defined as the percentage of unique peptides that are encoded by 2 or more oligonucleotides). The “coverage”, that is the percentage of recovered designed sequences, is provided at oligonucleotide level as well as peptide level. The total coverage of obtained by sequencing the naive phage libraries are shown alongside the predicted maximum coverage and error results, obtained by fitting the coverage vs. number of reads to a double hyperbolic curve.

**SI Table S2. Overview of protein domains and expression vectors used for protein expression.**

Details of the 40 expression constructs used with their UniProt accession numbers, gene name, domain name, construct start, construct end, the expression vector and the source of the construct.

**SI Table S3. PepTool annotated medium/high confidence ligands of the benchmarking set of bait proteins based on the HD2 P8 library screen.**

**SI Table S4. ProP-PD benchmarking set.**

Dataset of validated 466 motif instances for ProP-PD benchmarking compiled from the ELM database and structurally solved peptide-bait complexes from the Protein Data Bank (PDB). Each instance is mapped to a specific bait screened in the current analysis and is annotated with whether it is present in the HD2 library (*in library check*) and the number of overlapping peptides that are present in the HD2 library (*in library coverage*).

**SI Table S5. Overview of affinity data.**

K_D_, K_I_ and IC_50_ values generated in the study through FP direct binding or competition experiments.

**SI Table S6. Literature curated human WW domain-binding motifs.**

Dataset of 124 manually curated experimentally validated WW domain-binding motif instances collected from the literature.

**SI Table S7. Combined medium/high confidence data from selections against the HD2 P8 library, the HD2 P8 sub-libraries and the HD2 P3 library.**

Dataset of the *medium/high confidence* ligands selected across all screens. Data is available as a single combined sheet and as separate sheets for each sub-library. Data is organized on the peptide level allowing the reproducibly selected peptides for each bait to be investigated. Table includes filtered subsets of the combined dataset including ligands that are known interactors, in the dark proteome (Pharos *Tdark*), clinically relevant (Pharos *Tclin*), are co-localized (shared significant localization p-value < 0.05), or share significant biological processes (shared significant biological process p-value < 0.0005) or molecular function (shared significant molecular function p-value < 0.0005) GO-terms with a given bait.

**SI Table S8. Disease-related mutations in ProP-PD peptides.**

Dataset of the *medium/high confidence* ligands returned from the ProP-PD selections that overlap with disease-relevant mutations. Data is split into two sheets: (i) the complete dataset of disease-relevant mutations overlapping the selected ProP-PD peptides; and (ii) disease-relevant mutations overlapping matches to the defined positions in the ELM consensus for the bait and their 2 flanking residues within the selected ProP-PD peptides

**SI Table S9. Post-translational modifications in ProP-PD peptides.**

Dataset of the *medium/high confidence* ligands returned from the ProP-PD selections that overlap with PTMs. Data is split into two sheets: (i) the complete dataset of PTMs overlapping the selected ProP-PD peptides; and (ii) PTMs overlapping matches to the defined positions in the ELM consensus for the bait and their 2 flanking residues within the selected ProP-PD peptides

## ON-LINE METHODS

### Computational ProP-PD library design

#### Defining the ProP-PD search space

We defined the ProP-PD search space as the intrinsically disordered regions (IDRs), including loops in structured regions, of the human proteome accessible to intracellular proteins. A dataset of the 20,206 reviewed human proteins was retrieved from UniProt (release 2018_02) ^1^. Intracellular protein regions were defined by removing: (i) proteins with the keywords ‘Secreted’, unless they also had the keywords ‘Cytoplasm’ or ‘Nucleus’; and (ii) transmembrane regions and the extracellular regions of transmembrane proteins based on UniProt annotation. IDRs and large loops in structured regions of the human proteome were defined using 3 sources of data: (i) disorder state predictions using IUPred, (ii) surface accessibility from solved structures of the protein; and (iii) surface accessibility homology mapped from solved structures.

#### Defining the disordered regions of the human proteome

The disorder state predictions used IUPred to calculate per residue disorder propensity scores. Scores were calculated on the full length sequence of proteins from the UniProt dataset. In the cases where UniProt annotated chain and topology domains were available, the chain and topology domains of the protein were analyzed independently and these data were used. An IUPred disorder propensity score cut-off of 0.4 was applied to each residue of each protein resulting in binary accessible (disordered) or inaccessible residue classifications.

When a solved structure(s) of a protein was available, surface accessibility (SA) scores were calculated for the structure(s). The SA score for a residue was calculated as the proportion of the amino acid that is accessible to water molecules in the solved structure normalized by the maximum possible accessibility for that amino acid in a peptide chain (as defined for 5-residue peptides with a central query amino acid flanked by two glycine residues (GGXGG)). The SA score for a residue that is unresolved in the structure was set to 100% accessibility. For protein structures containing a multiprotein complex, regions that are less than 25 amino acids in length are discarded as they are unlikely to fold in the absence of a binding partner and chains with less than 10 intramolecular contacts per residue on average are not retained as this is a hallmark of a bound IDRs. When multiple structures are mapped to the same residue the median SA score for the residue is used. A SA cut-off of 33% is applied resulting in binary accessible or inaccessible residue classifications.

Homology mapped structures are defined by BLASTing the query protein against a database of PDB structure constructs and hits with an e-value cut-off of 10^-15^ and coverage cut-off of 85% of the structure are retained as homology mapped structures. The query protein and PDB structure constructs are aligned using local pairwise alignment with a BLOSUM62 matrix. SA scores are calculated for the PDB structure as described above and the SA scores were mapped using the pairwise alignment to the query protein. As above, a SA cut-off of 33% is applied resulting in binary accessible or inaccessible residue classifications.

The accessible/inaccessible categories to define IDRs and loops information were used in a hierarchy: direct or homology-mapped experimental accessibility data was used when available, otherwise disorder predictions were used (i.e. when experimental information is available it was used in place of predictions). The resulting binary accessible/inaccessible categories were smoothed to remove short regions of length 4 or less that are not consistent with the flanking region category. Regions of order in a disordered region that are less than 25 amino acids in length were defined as accessible and retained. Any 16-mer peptide window where at least 8 of the 16 amino acids were defined as accessible based on the rules above was retained as the ProP-PD search space.

#### Defining the peptides

The ProP-PD search space was tiled with peptides of length 16 amino acids overlapping by 12 amino acids. Cytoplasmic loops of length 8 or greater that are predicted as disordered were retained. All cysteines were replaced with alanine to avoid issues with unpaired cysteines in the phage coat proteins.

#### Design of the oligonucleotides

The designed peptide sequences were reverse translated into oligonucleotides by stochastically choosing codons to match the codon usage of *E. coli*. For special cases, i.e. when no overlapping peptide exists or the peptide was at a terminus, we created two distinct oligonucleotides for the peptides. The primers required for annealing in the construction of the phagemid library were added: (5’ CAGCCTCTTCATCTGGC and 3’ GGTGGAGGATCCGGAG). Finally, we redesigned oligonucleotides with a SmaI restriction site (GGGCCC or CCCGGG) or self-complementarity of greater than 7 contiguous nucleotides.

#### Defining HD2 sub-libraries

Peptides from the human proteins were split into 5 protein pools using Gene Ontology (GO) and UniProt Keyword annotation:

- *<u>Endomembrane:</u>* Mapping to GO terms “endomembrane system” or its descendants, or the UniProt Keywords “Endoplasmic reticulum membrane”, “Endoplasmic reticulum”, “Golgi apparatus membrane”, “Golgi apparatus”, “Golgi cisterna membrane”, “Golgi membrane”, “ER to Golgi transport vesicle membrane”, “Cytoplasmic vesicle membrane”, “Cytoplasmic vesicle”, “Early endosome membrane”, “Early endosome”, “Endosome membrane”, “Late endosome membrane”, “Late endosome” or “Recycling endosome membrane”.
- *<u>Nuclear:</u>* Mapping to GO term “nucleus” or “chromosome”, or their descendants, or the UniProt Keywords “Nucleus” or “Chromosome”.
- *<u>Cytoplasmic:</u>* Mapping to GO terms cytoplasm, mitochondrion, cytoskeleton, cilium or plasma membrane or their descendants, or the UniProt Keywords “Cytoplasm”, “Cell membrane”, “Membrane”.
- *<u>Extracellular:</u>* No *Cytoplasmic, Nuclear* or *Endomembrane* sub-library assignment. No transmembrane regions in the protein. Mapping to GO terms “extracellular region” or “extracellular region part” or their descendants, or the UniProt Keywords ‘‘extracellular space”, “extracellular exosome”, ‘‘extracellular region”, “extracellular exosome”, “exocyst”, “extracellular space”, “endoplasmic reticulum lumen” or “Endoplasmic reticulum lumen”.
- *<u>Other sub-library:</u>* Proteins without localization and therefore no sub-library assignment

Designed oligonucleotides were split into pools of 92,918 to match the size of the pools generated by the commercial provider. From these pools, we created 6 sub-libraries of different sizes:

- Cytoplasm (Cytoplasm and no Nucleus or Endomembrane System - 3 x 92.9 k pools),
- Nucleus (Nuclear and no Cytoplasm or Endomembrane System - 3 x 92.9 k pools),
- Endomembrane System (Endomembrane - 2 x 92.9 k pools),
- Nucleocytoplasmic shuttling (Nucleus and Cytoplasm or Endomembrane System - 4 x 92.9 k pools),
- Extracellular (Extracellular 1 x 92.9 k pools)
- Other (1 x 92.9 k pools).

Unused space on the chips for each sub-library was filled with redundant peptides encoded by distinct synonymous oligonucleotides. An interactive website to explore the full library design is available at http://slim.icr.ac.uk/phage_libraries/human/.

### Construction of the phage display libraries

The designed oligonucleotide libraries were obtained from CustomArray, and used to construct phage libraries displaying the encoded peptides on the P8 protein or the P3 protein following published protocols ^2^. Oligonucleotides from each sub-library pool were combined and used as template for 15 cycles of PCR amplification (denaturation at 98°C for 10 sec, annealing at 56°C for 15 sec and amplification at 72°C for 10 sec) using Phusion polymerase (Thermo Scientific) and primers complementary to the constant annealing regions flanking the designed library sequences. Remaining oligonucleotides and nucleotides were removed by *ExoI* (Thermo Scientific) treatment (HD2 P8) or using a nucleotide removal kit (Qiagen) (HD2 P3). The cleaned PCR products were phosphorylated using T4 polynucleotide kinase (Thermo Scientific) for 1 h at 37°C and annealed to phagemid ssDNA (90°C for 3 min, 50°C for 3 min and 20°C for 5 min). Later, dsDNA was synthesized using T7 DNA polymerase (Thermo Scientific) and T4 DNA ligase (Thermo Scientific) at 20°C for 16 h. The dsDNA generated from each sub-library pool was purified from a 1% agarose gel, eluted using ultrapure H20, and electroporated to *E.coli* SS320 (Lucigen) electrocompetent cells pre-infected with M13KO7 helper-phage (ThermoFisher) prepared as described elsewhere ^3^. The phage were allowed to propagate for 24 h in 2xYT (10 g yeast extract, 16 g tryptone, 5 g NaCl per L) medium. The phage was precipitated from the supernatant by the addition of 1/5^th^ volume of 20% PEG8000/2.5 M NaCl followed by centrifugation at 27,000 x g for 20 minutes. The phage pellets were dissolved in phosphate-buffered saline (PBS, 137 mM NaCl, 2.7 mM KCl, 95 mM Na_2_HPO_4_, 15 mM KH_2_PO_4_ pH 7.5). The sub-library phage titers were determined before pooling them into the final HD2 library. The resulting HD2 P8 library was reamplified and stored at −80°C.

### Next generation sequencing (NGS) of the phage datasets

Peptide-coding regions of the naive phage libraries or binding enriched phage pools (5 μL for 50 μL PCR reaction) were amplified and barcoded using a dual barcoding strategy ^4^ using Phusion High-Fidelity polymerase (Thermo Scientific) for 22 cycles ^2^. The PCR products were confirmed through 2% agarose gel electrophoresis stained with GelRed, and using a 50 bp marker (BioRad). The amount of the PCR products (25 μL) were normalized using 25 μL Mag-bind Total Pure NGS (Omega Bio-tek). The normalized amplicons were pooled (10 μL from each reaction). The resulting amplicon pool was further purified from a 2% agarose gel (QIAquick Gel extraction Kit Qiagen) with GelRed staining and eluted in TE (10 mM Tris-HCl, 1 mM EDTA. pH 7.5) buffer following the provided protocol, except dissolving the agarose gel by 30 min incubation at room temperature instead of vortexing. The dsDNA concentration of the purified amplicon pool was determined with Quant-iT PicoGreen dsDNA Assay Kit (Molecular probes by Life technologies) and analyzed by Illumina MiSeq v3 run, 1×150bp read setup, 20% PhiX by the NGS-NGI SciLifeLab facility, which returned an average of 33,000 reads per set of barcodes (on average 18 million reads per NGS run).

### Demultiplexing and processing of NGS data

NGS results processing was performed as described in detail in Ali *et al* ^2^. In brief: the results of ~500 pooled experiments, each one tagged with a unique combination of 5’ and 3’ barcodes, were demultiplexed and cleaned using in house custom Python scripts. Reads with an average quality score of 20 or more were retained, and their adapter regions and barcodes were determined allowing a maximum of 1 mismatch per adapter and/or barcode. Reads with ambiguous barcodes presenting a mutation that by allowing 1 mismatch can match them to more than 1 of the designed barcodes were excluded. The subset of high quality, unambiguously identified reads were trimmed by removing their adapter and barcode regions. The coding sequences were grouped into separated FASTA files where one file was produced for each barcode set. Finally, tables were built from each demultiplexed FASTA file where each unique trimmed oligonucleotide was translated to an amino acid sequence and NGS sequencing count.

### Library coverage and quality check

To assess the quality of synthesized and cloned phage libraries we analyzed the coverage percentage of the physical phage library compared to the designed phage library. Multiple aliquots for each naive phage library were sequenced through NGS and the total number of combined unique sequences matching the library design was used to calculate the “sequenced coverage”. For the estimation of the “maximum coverage”, each sequenced aliquot was picked in a random order and their cumulative contribution of new unique sequences versus the total number of contributed reads was assessed. The process of randomizing the picking order of sequenced aliquots was repeated 10 times and the maximum coverage was estimated by extrapolation to an infinite number of reads by non-linear regression to the following equation:

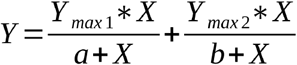

The “predicted maximum coverage” is the sum of Y_max1_ and Y_max2_, as determined by KaleidaGraph (Synergy Software) and the reported error is the propagated fitting error of this sum. As the HD2 P8 library was constructed from the HD2 sub-libraries, the contribution of each sub-library was also taken into account when calculating the sequenced and maximum coverage of the HD2 P8 library.

### Expression & purification of bait proteins for phage selections

*E.coli* BL21(DE3) gold bacteria (Agilent Technology) containing plasmids encoding 6-His-GST fusion proteins (**SI Table S2**) were grown in 100 mL 2xYT at 37°C and 200 rpm. For each protein, expression was induced with 1 mM isopropyl β-D-1-thiogalactopyranoside (IPTG) and was allowed to proceed for 4 h at 30°C. Bacteria was harvested for 10 min at 4,500 xg. The bacterial pellet was resuspended in lysis buffer (7.5 mL PBS supplemented with 1% Triton, 10 μg/mL DNaseI, 5 mM MgCl_2_, lysosome, and cOmplete™ EDTA-free Protease Inhibitor Cocktail (Roche)) and was incubated for 1h on ice. The suspension was sonicated to destroy remaining DNA and support the lysis, and the cell debris was pelleted by centrifugation (1h, 16,000 x g). Proteins were batch purified from the supernatant using GSH Sepharose 4 Fast Flow Media (Cytiva) following the manufacturer’s protocol. The protein concentration was estimated using a Nanodrop ND-1000 spectrometer and the purity was confirmed through SDS-PAGE.

### ProP-PD selections

Phage selections were performed following the previously described protocol ^2,5^. In brief, 10 μg GST-tagged bait protein or GST (total volume 100 μL in PBS) was immobilized in 96 well Flat-bottom Immunosorp MaxiSorp plates (Nunc, Roskilde, Denmark) for 18 h at 4°C. Wells were blocked with 200 μL 0.5% (w/v) BSA in PBS for 1 h, at 4° C. GST-coated wells were washed 4 times with 200 μL PT (PBS+0.05% (v/v) Tween20). The phage library (10^11^ phage in 100 μL PBS) was transferred to each of the GST-coated wells for removal of phage that bound non-specifically. After 1 h the library was transferred to protein coated wells. After 2 h of incubation unbound phage were removed by washing the wells with 5 x 200 μL PT. Bound phage were eluted with 100 μL log phase *E.coli* OmiMAX for 30 min at 37° C. Bacteria were hyperinfected by the addition of 10^9^ of M13KO7 helper phage per well for 45 min at 37° C. The bacteria (100 μL) were transferred in 1 mL 2xYT supplemented with 100 μg carbenicillin, 30 μg kanamycin, 0.3 mM IPTG and were incubated at 37°C for 18 h upon shaking. Phages were harvested by centrifugation at 2,000 x g for 10 min. Phage supernatants were transferred into a fresh 96-well plate, pH adjusted by the addition of 1/10 volume 10x PBS and heat inactivated for 10 min at 65° C. The resulting amplified phage pools were used as in-phage for the next day of selection. To enrich for binding phage, this procedure was repeated in total in four rounds of selections.

To evaluate the enrichment of binding phage, pooled phage ELISA was performed in 384 well Flat-bottom Immunosorp MaxiSorp plates (Nunc, Roskilde, Denmark). For each round of selections bait protein and GST control were coated (5 μg in 50 μL/well) for 18h at 4° C. The wells were blocked with 100 μL 0.5% BSA in PBS, for 1 h, at 4° C. Phage (50 μL) from the different rounds of selections were allowed to bind to the bait protein or GST coated wells. After 1 h unbound phage was washed away with 4x 100 μL PT. Bound phage was detected using 50 μL M13 HRP-conjugated M13 bacteriophage antibody (Sino Biological Inc; 1:5000 diluted in PT, 0.05% BSA) for 1 h, at 4° C. The wells were washed with 4x 100 μL PT and once with 100 μL 1x PBS. TMB substrate (40 μL, Seracare KPL) was added to detect the bound antibody. The enzymatic reaction was stopped by the addition of 40 μL 0.6 M H_2_SO_4_. The absorbance at 450 nm was determined with an iD5 (Molecular Devices).

### ProP-PD peptide analysis

#### Peptide Processing

Peptide data was created for each ProP-PD selection as described in “Demultiplexing and processing of NGS data”. Peptides observed only a single time in the NGS sequencing data were discarded. The NGS sequencing counts for each selection day were normalized by dividing the peptide sequencing counts by the sum of the sequencing counts for all peptides in the selection to create the *normalized peptide sequencing count*. Peptides for each selection day for a replicate were pooled and the mean *normalized peptide sequencing count* for each replicate was calculated for each peptide.

#### PepTools peptide mapping and annotation web server

We developed PepTools, a novel peptide annotation tool, to analyse peptide data from experimental motif discovery methods such as ProP-PD. PepTool expands on the SLiMSearch and PSSMSearch motif discovery framework ^6,7^ to map peptides to a query proteome and annotate them with structural, evolutionary, functional, genomic and proteomic data. The PepTools framework is available as a web server that is freely accessible at http://slim.icr.ac.uk/tools/peptools/.

PepTools has a range of functions and features to quickly pinpoint putative biologically relevant motif instances in a list of peptides:

- *Previously validated instances*: overlap with previously validated motif instances retrieved from the ELM database.
- *Previously validated interactions with bait protein/domain:* evidence of an interaction of the peptide-containing protein with a bait protein or a protein containing a bait domain.
- *Shared annotation with bait protein/domain:* shared functional annotations or localization of the peptide-containing protein with a bait protein or a protein containing a bait domain (calculated as described previously ^9^).
- *Accessibility information*: accessibility information from annotations (topology, domains), predictions (intrinsic disorder, transmembrane regions) and experimental sources (structure-derived surface accessibility).
- *Consensus/PSSM annotation and filtering:* peptides can be scanned with motif consensuses to highlight peptides with motif matches and peptides can be scored and ranked with a user-defined PSSM. Peptides can also be filtered using the motif consensus or peptide PSSM scores.
- *Key specificity determinant residue annotation:* residue-specific annotation such as SNPs, PTMs and mutagenesis are highlighted if they overlap “key” residues based on a user-defined motif consensus.
- *Enriched motif specificity determinants:* Peptides can be analyzed for enriched motifs using the SLiMFinder motif discovery tool ^22^. The motif specificity determinants of the enriched motifs can be visualized with a heatmap or sequence logo.
- *User annotation:* External peptide related data can be added to the input resulting in sortable extra columns on the results page.
- *Extensive peptide filtering functionality:* ontology, interaction and localization information allow the use of *a priori* knowledge of a motif’s biological context for peptide prioritisation.
- *Masked amino acid:* specific amino acids can be masked to allow peptides with experimentally required substitutions to be mapped correctly (for example, cysteine to alanine substitutions in a ProP-PD library).
- A detailed description of PepTools functionality is available on the PepTools help page at http://slim.icr.ac.uk/tools/peptools/help
- The filtering, enrichment and annotation functionality is as described for the SLiMSearch and PSSMSearch ^6,7^.

Key sources of data used by PepTools for the ProP-PD analysis include (i) protein data (UniProt); (ii) localization data (UniProt ^1^ and Gene Ontology ^8^); (iii) accessibility data (structure data retrieved from PDB processed using DSSP and IUPred calculated disorder scores ^9,10^); (iv) domain data (retrieved from Pfam ^11^); (v) interaction data (retrieved from IntAct ^12^, HIPPIE ^13^, BioPlex ^14,15^ and HuRI ^16^); (vi) post-translational modifications (retrieved from the UniProt ^1^, phospho.ELM ^17^ and PhosphoSitePlus ^18^ database and a large phosphoproteomics dataset from Ochoa *et al*^19^); and (vii) disease-relevant SNPs (data from gnomAD, dbSNP, COSMIC curated, TOPMed, NCI-TCGA COSMIC, NCI-TCGA, ExAC, Ensembl, ESP, ClinVar, 1000Genomes retrieved from the EBI API ^20^ and UniProt Human polymorphisms and disease mutations ^1^).

#### PepTools peptide mapping and annotation

Selected peptides matching the designed peptide length (16 amino acids) were mapped to the human proteome using PepTools. Alanine residues were permitted to be variable in the PepTools peptide mapping to allow the remapping of cysteine residues converted to alanine in the ProP-PD library design. Peptides were annotated with structural, evolutionary, functional, genomic and proteomic data. For each bait, the peptides for each replicate were compared to define replicated peptides and overlapping peptides. Overlapping peptides were mapped using their protein mapping (rather than sequence) and the peptides defined as overlapping can be present in different replicates for the same bait. For the benchmarking analyses, to remove biases caused by peptides that mapped to two or more closely related paralogues, peptides were mapped to a single primary paralogue as defined by the UniProt Reference Clusters (UniRef) clusters.

#### Motif enrichment, specificity determinant score and motif-containing peptides

The replicated peptides and overlapping regions of overlapping peptides for each bait were analyzed for enriched motifs using the SLiMFinder motif discovery tool ^6^. SLiMFinder was run using *minic* (Minimum information content for returned motifs) of 1.1, *equiv* (list of groupings of physicochemically similar residues to use for ambiguous positions) of AGS,ILMV,IVMLFAP,IVMLFWHYA,FYWH,KRH,ST,STNQ,DEST and *maxwild* (the maximum number of consecutive wildcard positions to allow) of 5. The amino acid frequencies of the input peptides were used as background amino acid frequencies. The top-ranked SLiMFinder motif from the motif enrichment step was used to align the motif-containing peptides and a PSI-BLAST PSSM was built from the aligned peptides using the PSSMSearch motif discovery tool ^7^. The PSSM was used to calculate a specificity determinant score, a metric to define the similarity of a peptide to the enriched specificity determinants, for the selected peptides for a bait. The PSSM specificity determinant score was calculated using the probabilistic scoring method of the PSSMSearch motif discovery tool on a background model obtained from scanning the human proteome with the reversed variant of the PSSM.

#### Specificity determinant visualization

Specificity determinants were visualized as sequence logos created using the PSSM visualization software of the PSSMSearch tool. The logos display relative binomial amino acid frequencies calculated as *log^-10^* of the binomial probability (*prob^aa^ = binomial(k,n,p)* where k is the observed residue count at each position for a residue, *n* is the number of the instances of motifs and *p* is the background frequency of the residue in the disordered regions of the human proteome). In the case of the multiple motifs generated for the KPNA4 binding peptides, peptides were first separated into different cohorts following the previously established classification system ^21^.

### Selection amino acid bias analysis

Amino acid frequencies were calculated for the human proteome (UniProt reviewed human proteins (release 2018_02)); the predicted disordered regions of the human proteome (IUPred score > 0.4 for the UniProt reviewed human proteins); the HD2 library design; the binding enriched phage pools from selections against HD2 P8 library; and the binding enriched phage pools from selections against combinatorial P8 library. The relative amino acid frequencies were calculated for the HD2 P8 and the combinatorial peptide phage display versus the amino acid frequencies of predicted disordered regions of the human proteome.

### Screen quality checks

#### Selection replicates benchmarking

All ProP-PD selections were compared in a pairwise manner and the proportion of selected peptides (i) shared between the selections, or (ii) overlapping between the selections were calculated. Each ProP-PD screen pairwise comparison was classified as replicate selections for the same bait, the same control bait and different bait proteins. The proportion of replicated and overlapping peptides in replicate selections was then compared to control and non-replicate selections.

#### Enriched consensus benchmarking

Three consensus-based metrics were calculated for each screened bait: (i) the enrichment of the expected ELM consensus in the peptides selected for the bait, (ii) the enrichment of a *de novo* consensus defined by SLiMFinder in the peptides selected for the bait, and (iii) the similarity of the *de novo* SLiMFinder consensus to the expected ELM consensus.

The ELM defined class(es) were curated for each bait. The enrichment of each ELM class consensus(es) was calculated in the set of peptides returned from each bait using the binomial probability (*prob^aa^* = *binomial(k,n,p)* where k is the number of selected peptides for the bait that match the consensus(es), *n* is the number of the peptides for the bait and *p* is the frequency peptides matching the consensus(es) in the whole HD2 library). The consensus enrichment for the correct consensus-bait pairs was then compared to all other consensus-bait pairs.

For each bait, the replicated peptides and overlapping regions of overlapping peptides were analyzed for enriched motifs using the SLiMFinder motif discovery tool ^22^ as defined in *“Motif enrichment, specificity determinant score and motif-containing peptides’“*. The most significant returned consensus above a p-value cut-off of 0.001 was defined as *de novo* SLiMFinder-defined enriched motifs for the bait. The *de novo* SLiMFinder-defined enriched motif was then compared with the correct ELM-defined consensus(es) for the bait and against the consensus for all other ELM classes using the CompariMotif software ^22^.

### ProP-PD motif benchmarking

#### ProP-PD motif benchmarking datasets

We defined the *ProP-PD motif benchmarking dataset* to test the ability of the ProP-PD method to discover motifs. The dataset was created from 466 motif instances that were previously validated as binding to the 40 bait proteins tested in the study (**SI Table S4**). Of these, 337 were covered by one or more peptides in the HD2 library and bound to one of the 35 non-control baits. The motifs were compiled from the ELM database ^23^ and structurally solved peptide-bait complexes from the PDB ^24^. For the ELM instances, each bait was annotated to an ELM class or classes and all motif instances for that class were defined as validated binders for the bait. The PDB instances were collected by retrieving structures of protein complexes that contain the bait from PDB and computationally parsing peptides bound to the domain used in the ProP-PD screens. A further manually curated WW domain-binding motif dataset was also collected. A list of 124 PPxY motif instances experimentally validated to bind to WW domains was collected from the literature and used to create a benchmarking dataset to evaluate ProP-PD selected peptides for the NEDD4 and YAP1 WW domain screens (**SI Table S5**).

#### Validated motif benchmarking

The ProP-PD selection data was benchmarked on the *ProP-PD motif benchmarking datasets*. ProP-PD selected peptides that overlap with motif instances in the *ProP-PD motif benchmarking datasets* were compared to all other selected peptides. Four peptide metrics were compared: replicated peptides (the number of replicates that the peptides are observed in), overlapping peptides (the number of distinct peptides overlapping the peptide across all replicates), specificity determinant match (the SLiMFinder-derived PSSM match p-value) and normalized peptide count (the mean normalized peptide count for the peptide across the NGS counts of the replicates). The predictive power was calculated for each metric, defined by the area under the ROC curve (AUC) and Mann-Whitney-Wilcoxon two-sided test with Bonferroni correction p-value.

### Peptide consensus confidence assignment

Optimal cut-offs for each of the peptide metrics were calculated using Youden’s J statistic to maximize the true positive rate and minimize the false positive rate: (i) replicated peptide (the peptide is observed in two or more replicates); (ii) overlapped peptide (the peptide has one or more overlapping peptides in any replicate); (iii) specificity determinant match (the peptide has a SLiMFinder-derived PSSM match with a p-value < 0.0001); and (iv) high normalized peptide count (the peptide has a mean normalized peptide count > 0.0005). The four binary confidence criteria were combined for each peptide to create a single metric ‘Confidence level’ which has four categories (‘High’, ‘Medium’, ‘Low’ and ‘Filtered’). The ‘Confidence level’ of ‘High’ is assigned to instances matching all four confidence criteria, ‘Medium’ is assigned to instances matching two or three of the criteria and ‘Low’ is assigned to instances matching one metric criterion. Peptides that do not match any of the metric criteria are defined as ‘Filtered’ and discarded from the results.

### ProP-PD interaction benchmarking

We defined the *ProP-PD interaction benchmarking dataset* from the 302 interactions annotated for the 337 motif instances in the *ProP-PD motif benchmarking dataset* (**SI Table S4).** The 302 motif-mediated interactions were cross-referenced against high confidence ProP-PD interactions, the high/medium confidence ProP-PD interactions, the AP-MS-derived BioPlex 3.0 interaction dataset ^14^ and the Y2H-derived HuRI interaction dataset ^16^. Next, the overlap of the interactions from the high/medium confidence ProP-PD interactions, the BioPlex 3.0 interaction dataset and the Y2H-derived HuRI interaction dataset for interaction in the *ProP-PD interaction benchmarking dataset* was calculated. Finally, the overlap of the high/medium confidence ProP-PD interactions with the integrated human PPI dataset from HIPPIE was calculated ^13^.

### Comparison of results from HD2 P8 selections to selections against the sub-libraries and HD2 P3

Pairwise comparison between the high/medium confidence peptides generated from selections against the HD2 P8 library and the rest of the libraries (HD2 P3 and HD2 P8 sub-libraries) was made to evaluate how the results of selections against each distinct library compares to the results from selections against the HD2 P8 library. The comparison measured the extent to which selections against each library discovered known instances belonging to the *ProP-PD motif benchmarking dataset* (SI **Table S4**; http://slim.icr.ac.uk/data/proppd_hd2_pilot). The performance of each library was defined by two values, Recall and Precision. The recall was defined by the number of previously validated instances from the benchmarking set rediscovered in the high/medium set of ligands generated by selections against each library. The same benchmarking set was used to measure recall for HD2 P8 and the compared library. The precision was defined by the number of previously validated instances from the *ProP-PD benchmarking dataset* found in the medium/high confidence set of ligands in comparison to the total number of medium/high confidence ligands.

### Purification of KEAP1 Kelch, KPNB1 HEAT and MDM2 SWIB for affinity measurements

The protein coding regions (**SI Table S2**) were subcloned using the *EcoRI* and *NcoR1* restriction site into in pETM41 (EMBL) for the expression of 6-His-MBP-tagged proteins or in the pETM33 (EMBL) vector. Protein expression was induced at OD600 0.8 with 1 mM IPTG, and allowed to proceed at 18°C for 20 h. The bacteria were harvested and lysed under the same conditions as described above. The supernatant was batch purified using the IMAC technique with Ni Sepharose 6 Fast Flow (Cytiva). After 1 h incubation with the gel slurry, the gel was transferred into a column. The beads were washed with wash buffer (20 mM NaPO_4_, 0.5 M NaCl, 30 mM imidazole pH 7.4) and eluted with elution buffer (20 mM NaPO_4_ 0.5 M, NaCl 500 mM imidazole, pH 7.4). The proteins were further purified as follows:

- 14-3-3 SFN: The protein was not eluted from the IMAC column. Instead, after washing unbound protein away, His-tagged HRV3C protease was added and incubated for 16 h in 20 mM NaPO_4_, 0.5 M NaCl, pH 7.4. The cleaved protein was eluted with the same buffer. The cleaved protein was dialyzed into 50 mM KPO_4_ pH7.5 for 16 h.
- TLN1 PTB was cleaved with His-tagged HRV3C protease while dialysing into 50 mM KPO_4_ pH 7.5 for 16 h 4°C. The cleaved protein was applied on a Ni^2+^ IMAC gel and incubated for 1 h at 4°C to remove the tag. The protein was once more dialyzed in 50 mM KPO_4_ pH 7.5 for 16 h.
- MDM2 SWIB: The protein was cleaved from the His-GST tag using His-tagged HRV3C protease and then further purified with a S100 HiPrep 16/60 Sephacryl gel filtration using 150 mM NaCl, 20 mM NaPO_4_ pH 7.4. After gel filtration the protein was dialyzed into 50 mM NaPO_4_ pH 7.5 for 16 h.
- KPNB1 HEAT: The eluted protein was cleaved with His-tagged TEV protease while dialysing in 150 mM NaCl, 50 mM Tris, 0. 5 mM EDTA, 1 mM 1,4-Dithiothreitol (DTT) pH 8.0 for 16 h. The cleaved tag and the protease were removed through a reverse Ni^2+^ IMAC. The protein was dialyzed in 50 mM KPO_4_ pH 7.5, 1 mM DTT for 16 h.
- KEAP1 Kelch: The MBP-tagged domain was dialyzed into 50 mM KPO_4_ pH 7.5 for 16 h.
- KPNA4 ARM: The MBP-tagged domain was further purified through size exclusion chromatography using a Sephacryl S300 high resolution 26/600 and 50 mM KPO_4_ pH 7.5 as running buffer.

### Fluorescence polarization

FP affinity measurements were carried out with an iD5 multi detection plate reader (Molecular Devices) using Corning assay 96 well half area black Flat-bottom Non-binding surface plates (Corning, USA #3993). The settings were 485 nm Excitation and 535 nm for emission at a reading height of 1.76 mm and total volume of 50 μL. Peptides were obtained from GeneCust (France) at >95% purity. Unlabeled peptides were dissolved in 50 mM KPO_4_ or 50 mM NaPO_4_, pH 7.5. FITC-labeled peptides were dissolved in dimethyl sulfoxide (DMSO). Protein for saturation experiments, or peptides for the displacement experiments, were arrayed in serial dilution in 50 mM KPO_4_ pH 7.5 in 25 μL, followed by addition of 25 μL of a master mix. For saturation binding experiments, the master mix contained 2 mM DTT and 10 nM FITC-labeled peptide in 50 mM KPO_4_ pH 7.5 or 50 mM NaPO_4_ pH 7.5. For competition experiments, the master mix was supplemented with the protein of interest at a concentration of 4 times the K_D_ value. Data was analyzed with GraphPad Prism version 9.0.0 for MacOS (GraphPad Software, San Diego, California USA, www.graphpad.com). For direct binding, we used a quadratic equation with *pept* indicating the fixed probe peptide concentrations, *X* indicating the protein concentration, the constant *A* being the signal amplitude divided by probe peptide concentration, and B is the plateau value:

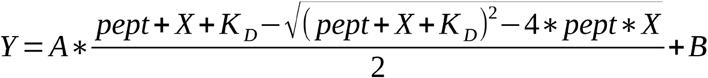

For the FP competition experiments data was fitted to a sigmoidal dose-response (variable response; GraphPad Prism).

### Cloning & mutagenesis for analysis of nuclear localization signals

For the cell-based NLS experiment using peptides, a pEGFP-C1 vector was modified to contain two additional eGFP genes spaced by a linker region on the 3’ end of the vector’s multiple cloning site (**SI Figure S7**). The modified vector is here called a “tri-GFP vector”. DNA strands for tandem peptides to be tested for NLS function were constructed by PCR using overlapping primers that coded for the peptide sequences and also contained overlapping regions for the MCS in the tri-GFP vector. The PCR products were then cloned into the tri-GFP vector using Gibson Assembly^®^ (New England Biolabs) following the manufacturer’s recommendations. Obtained clones were verified by Sanger sequencing.

NLS experiments using full-length target proteins were carried out using mCherry (Addgene plasmid #54563) tagged proteins. Briefly, DNA strands of full-length proteins were amplified using primers containing 5’ overhangs matching MCS of the target vector. The PCR products were later cloned into the mCherry2-C1 vector using Gibson Assembly^®^ (New England Biolabs) following the manufacturer’s recommendations. The putative NLS site in each construct was mutated by site directed mutagenesis to confirm the functional NLS sequence. All obtained clones were verified by Sanger sequencing.

### Cell culture and samples preparation

HEK293 cells were obtained from Sigma (Cat. 85120602) and cultured using DMEM with GlutaMAX™ Supplement (Gibco™) supplemented with 10% FBS (Gibco™) and Non-Essential Amino Acids Solution (NEAA, Gibco™). Cultures were maintained at 37 °C with 5% CO_2_ in humidified chambers and were routinely checked for mycoplasma contamination. For NLS experiments, cells were transfected using FuGENE^®^ HD (Promega) according to manufacturer’s instructions and using highly purified DNA samples. Cells were then grown in 8-well chamber slides. Following fused GFP or mCherry expression for 36 hours post-transfection, cells were washed with ice-cold PBS and were fixed using Image-iT™ Fixative Solution containing 4% formaldehyde (Invitrogen™) for 15 minutes on ice. Cells were washed 3 times with PBS each for 5 minutes at room temperature. Slides were dried and mounted with ProLong™ Glass Antifade Mountant with NucBlue™ Stain (Invitrogen™).

### Microscope image acquisition and processing

Images were acquired by Zeiss imager Z2 microscope using C11440 camera (Hamamatsu) and 40x oil objective lens (N.A. 1.4) using Zen software (V3.2, blue edition). HXP 120 V light source unit was used for sample excitations using fixed light intensities across all samples and images collected using appropriate filter sets. Merged images of 8-bit depth were exported and processed in ImageJ, where color brightness was adjusted for each channel homogeneously for all the images.

### Disease-relevant SNPs analysis

PepTools SNP annotation of the high/medium confidence peptides was analyzed. Data were filtered to create a “disease-relevant” SNP dataset based on clinical significance annotation (“Pathogenic”, “Likely Pathogenic”, “Disease”, “Risk factor”, “Association” “Protective”, “Drug response”, “Affects”). These “disease-relevant” SNPs were mapped to key specificity determinant residues (based on the defined positions in the ELM consensus for the given bait) and the two flanking residues. The output was then used to build a PPI network for each bait (**S1 Table S8**). In total, we constructed 16 networks, where each bait is binding at least to one peptide with a mutation. A network composed of only peptides with disease-associated mutations affecting motif encoding residues was also built and visualized using Cytoscape (**SI Fig. S8**).

### Phosphorylation analysis

PepTools phosphorylation site annotation of the high/medium confidence peptides was analyzed (**SI Table S9**). Phosphorylation sites were mapped to key specificity determinant residues (based on the defined positions in the ELM consensus for the given bait) and the two flanking residues. Phosphosite information was added to the disease-associated mutations PPI networks (**SI Fig. S8**).

### Plots and visualization

Graphs were created using the Matplotlib ^25^ and Seaborn ^26^ libraries in Python 3 ^27^, or with ggplot2 library ^28^ in the R scripting language ^29^. Structure figures were created with PyMOL ^30^. All networks were visualized using Cytoscape ^31^.

