## Supplemental Figures 1-8 for "Proteome-scale amino-acid resolution footprinting of protein-binding sites in the intrinsically disordered regions of the human proteome"

### SI Fig. S1

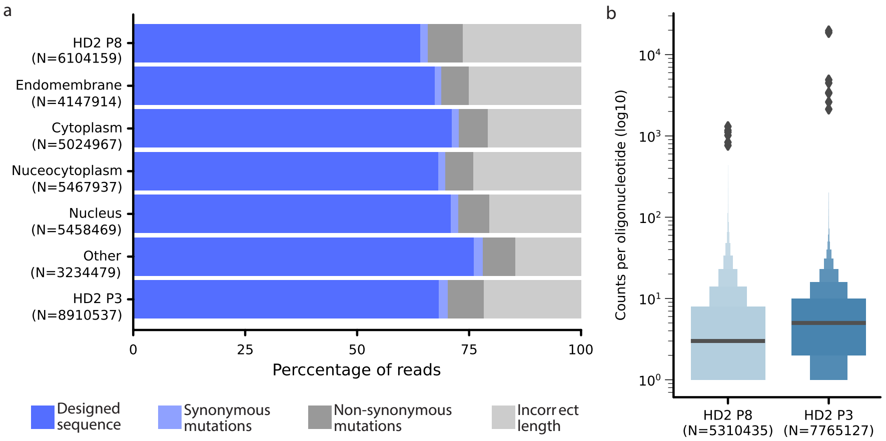

a
HD2 P8
(N=6104159)
Endomembrane
(N=4147914)
Cytoplasm
(N=5024967)
Nuceocytoplasm
(N=5467937)
Nucleus
(N=5458469)
Other
(N=3234479)
HD2 P3
(N=8910537)
0
25
50
75
100
Perccentage of reads
Designed sequence
Synonymous mutations
Non-synonymous mutations
Incorrect length
b
Counts per oligonucleotide (log10)
$10^4$
$10^3$
$10^2$
$10^1$
$10^0$
HD2 P8
(N=5310435)
HD2 P3
(N=7765127)

SI Fig. S2.

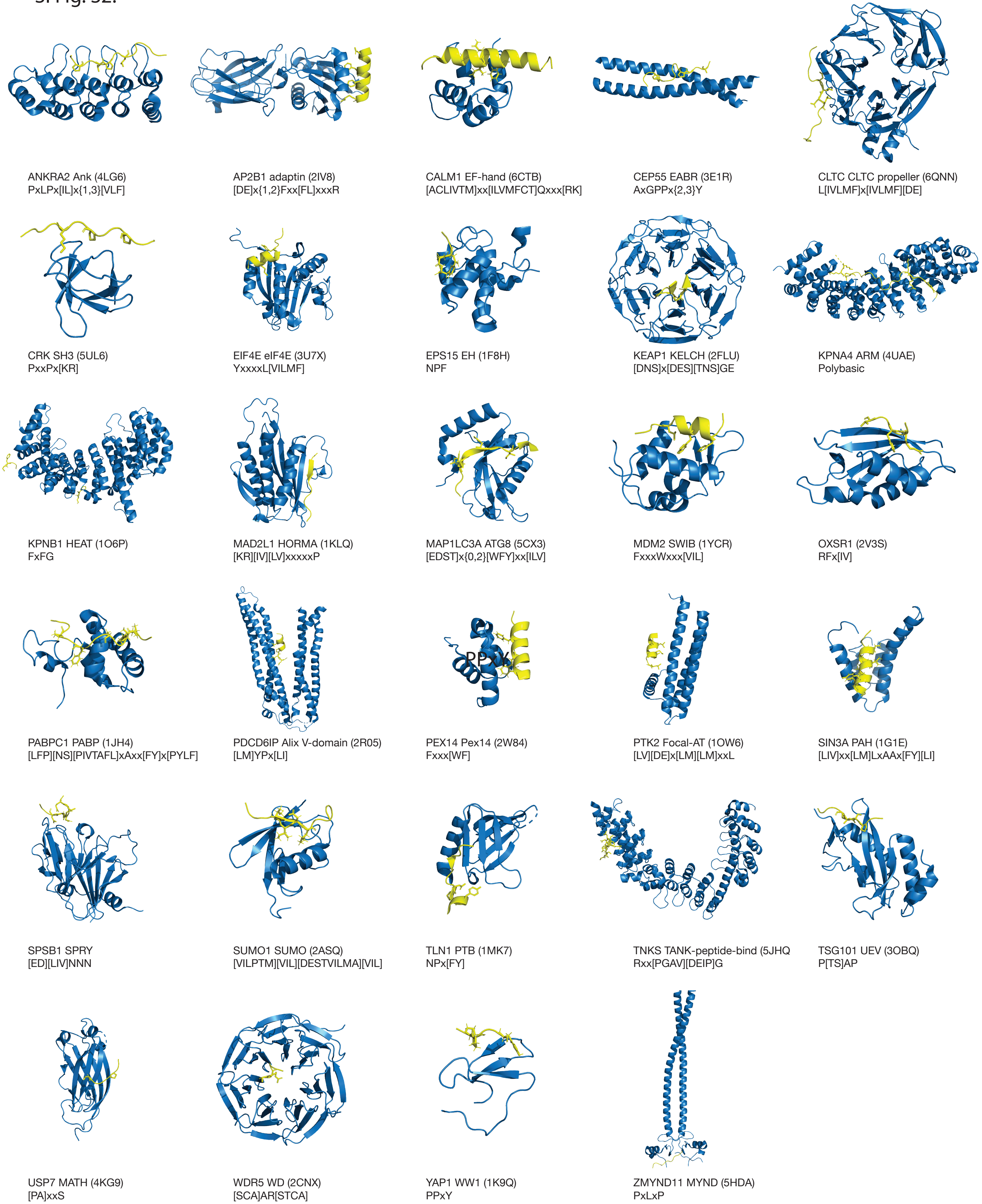
ANKRA2 Ank (4LG6)
PxLPx[IL]x{1,3}[VLF]
AP2B1 adaptin (2IV8)
[DE]x{1,2}Fxx[FL]xxxR
CALM1 EF-hand (6CTB)
[ACLIVTM]xx[ILVMFCT]Qxxx[RK]
CEP55 EABR (3E1R)
AxGPPx{2,3}Y
CLTC CLTC propeller (6QNN)
L[IVLMF]x[IVLMF][DE]
CRK SH3 (5UL6)
PxxPx[KR]
EIF4E eIF4E (3U7X)
YxxxxL[VILMF]
EPS15 EH (1F8H)
NPF
KEAP1 KELCH (2FLU)
[DNS]x[DES][TNS]GE
KPNA4 ARM (4UAE)
Polybasic
KPNB1 HEAT (1O6P)
FxFG
MAD2L1 HORMA (1KLQ)
[KR][IV][LV]xxxxxP
MAP1LC3A ATG8 (5CX3)
[EDST]x{0,2}[WFY]xx[ILV]
MDM2 SWIB (1YCR)
FxxxWxxx[VIL]
OXSR1 (2V3S)
RFx[IV]
PABPC1 PABP (1JH4)
[LFP][NS][PIVTAFL]xAxx[FY]x[PYLF]
PDCD6IP Alix V-domain (2R05)
[LM]YPx[LI]
PPxY
PEX14 Pex14 (2W84)
Fxxx[WF]
PTK2 Focal-AT (1OW6)
[LV][DE]x[LM][LM]xxL
SIN3A PAH (1G1E)
[LIV]xx[LM]LxAAx[FY][LI]
SPSB1 SPRY
[ED][LIV]NNN
SUMO1 SUMO (2ASQ)
[VILPTM][VIL][DESTVILMA][VIL]
TLN1 PTB (1MK7)
NPx[FY]
TNKS TANK-peptide-bind (5JHQ
Rxx[PGAV][DEIP]G
TSG101 UEV (3OBQ)
P[TS]AP
USP7 MATH (4KG9)
[PA]xxS
WDR5 WD (2CNX)
[SCA]AR[STCA]
YAP1 WW1 (1K9Q)
PPxY
ZMYND11 MYND (5HDA)
PxLxP

SI Fig. S3

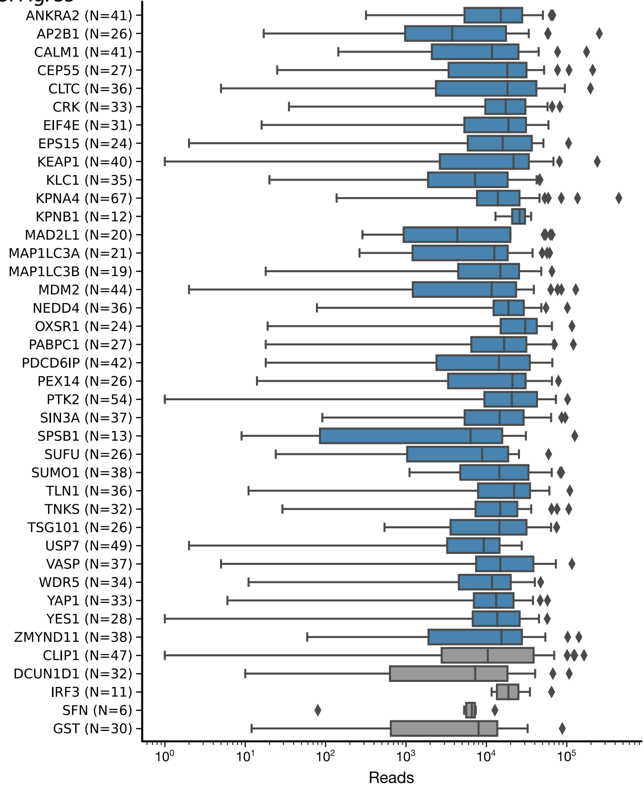
ANKRA2 (N=41)
AP2B1 (N=26)
CALM1 (N=41)
CEP55 (N=27)
CLTC (N=36)
CRK (N=33)
EIF4E (N=31)
EPS15 (N=24)
KEAP1 (N=40)
KLC1 (N=35)
KPNA4 (N=67)
KPNB1 (N=12)
MAD2L1 (N=20)
MAP1LC3A (N=21)
MAP1LC3B (N=19)
MDM2 (N=44)
NEDD4 (N=36)
OXSR1 (N=24)
PABPC1 (N=27)
PDCD6IP (N=42)
PEX14 (N=26)
PTK2 (N=54)
SIN3A (N=37)
SPSB1 (N=13)
SUFU (N=26)
SUMO1 (N=38)
TLN1 (N=36)
TNKS (N=32)
TSG101 (N=26)
USP7 (N=49)
VASP (N=37)
WDR5 (N=34)
YAP1 (N=33)
YES1 (N=28)
ZMYND11 (N=38)
CLIP1 (N=47)
DCUN1D1 (N=32)
IRF3 (N=11)
SFN (N=6)
GST (N=30)
$10^0$
$10^1$
$10^2$
$10^3$
$10^4$
$10^5$
Reads

SI Fig. S4

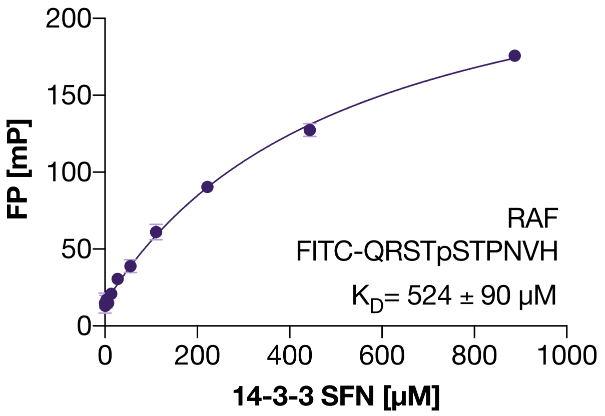
FP [mP]
14-3-3 SFN [µM]
RAF
FITC-QRSTpSTPNVH
$K_D$= 524 ± 90 µM

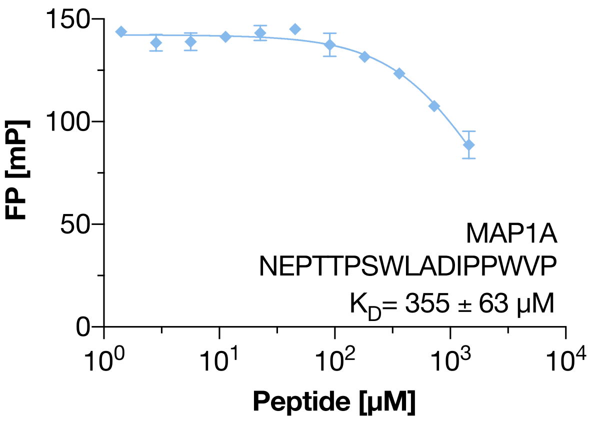
FP [mP]
Peptide [µM]
MAP1A
NEPTTPSWLADIPPWVP
$K_D$= 355 ± 63 µM

SI Fig. S5

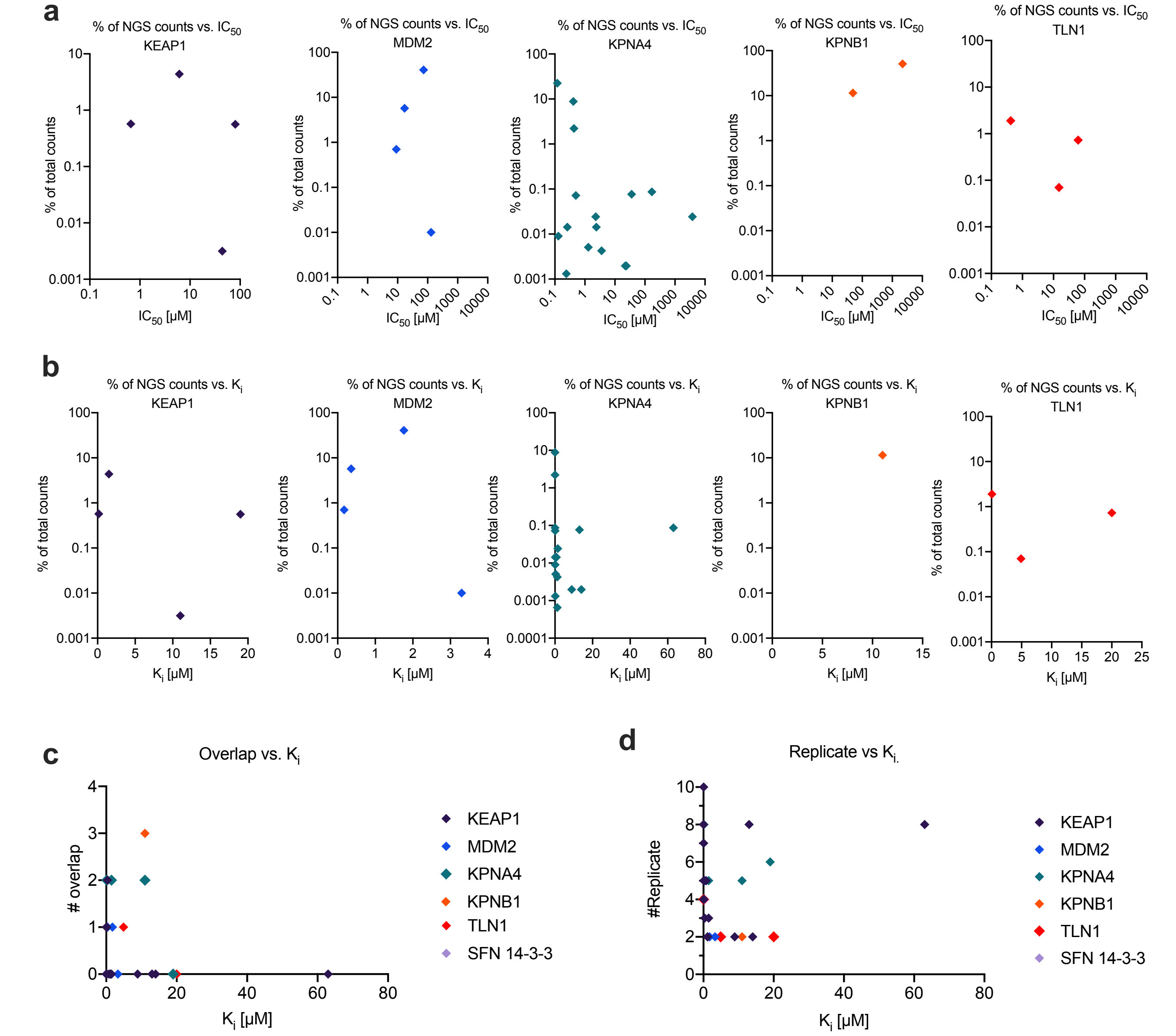
a
% of NGS counts vs. IC50
KEAP1
MDM2
KPNA4
KPNB1
TLN1
% of total counts
IC50 [μM]
b
% of NGS counts vs. Ki
Ki [μM]
c
Overlap vs. Ki
### overlap
d
Replicate vs Ki.
#Replicate
KEAP1
MDM2
KPNA4
KPNB1
TLN1
SFN 14-3-3

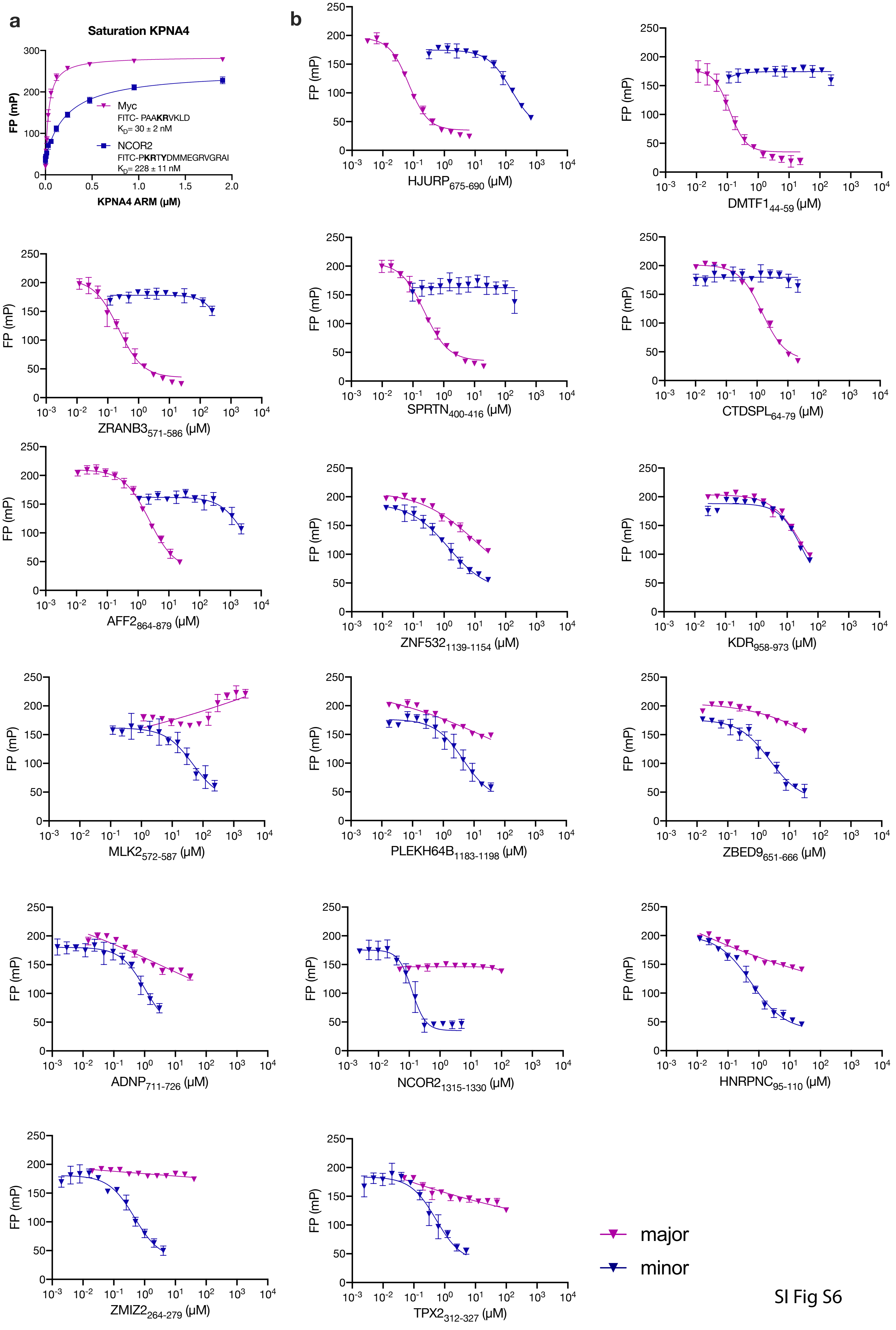
a
Saturation KPNA4
FP (mP)
KPNA4 ARM (µM)
Myc
FITC- PAAKRVKLD
KD= 30 ± 2 nM
NCOR2
FITC-PKRTYDMMEGRVGRAI
KD= 228 ± 11 nM
b
FP (mP)
HJURP675-690 (µM)
DMTF144-59 (µM)
ZRANB3571-586 (µM)
SPRTN400-416 (µM)
CTDSPL64-79 (µM)
AFF2864-879 (µM)
ZNF5321139-1154 (µM)
KDR958-973 (µM)
MLK2572-587 (µM)
PLEKH64B1183-1198 (µM)
ZBED9651-666 (µM)
ADNP711-726 (µM)
NCOR21315-1330 (µM)
HNRPNC95-110 (µM)
ZMIZ2264-279 (µM)
TPX2312-327 (µM)
major
minor

SI Fig S6

SI Fig S8

**A**

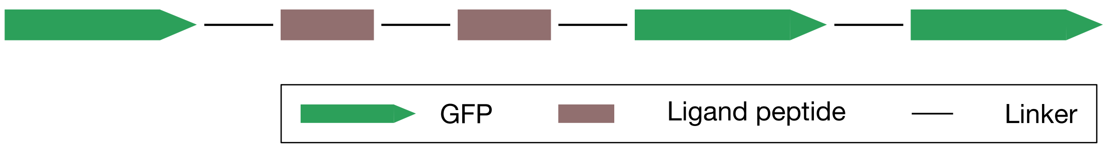


**B**

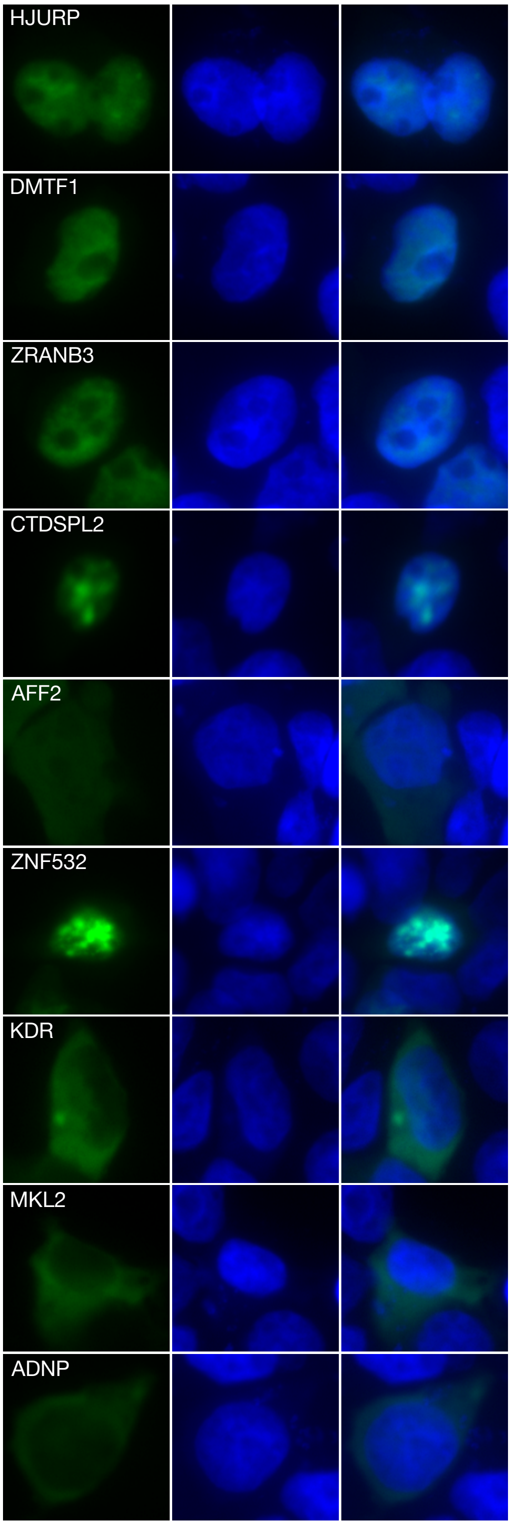


**C**

| Protein | Peptide sequence | Nuc / Cyto |
|---|---|---|
| HJURP | 675- DHQFPAKRPRLSEPQG -690 | Nuc |
| DMTF1 | 44- EDSIEPPHKRLALSSE -59 | Nuc |
| ZRANB3 | 571- VEPETKRLKLAASEDH -586 | Nuc |
| SPRTN | 401- EDTFPNKRPRLEDKTV -416 | Nuc |
| CTDSPL2 | 64- ERENPSKRSRIERDID -79 | Nuc |
| AFF2 | 864- IPEKKQRLEEATTICL -879 | Cyto |
| ZNF532 | 1139- PSPKRKLEEPVLEFRP -1154 | Nuc |
| KDR | 958- DLKRRLDSITSSQSSA -973 | Cyto |
| MKL2 | 572- ELKRKLEQEQKLVEVL -587 | Cyto |
| PLEKHG4B | 1183- PFKRPHSTISDSSTSS -1198 | Nuc |
| ZBED9 | 651- STMKRKHDEIQRSLPV -666 | Nuc |
| ADNP | 711- SLAPVKRTYEQMEFPL -726 | Cyto |
| NCOR2 | 1315- PKRTYDMMEGRVGRAI -1330 | Nuc |
| HNRNPC | 95- AGVKRSAAEMYGSVTE -110 | Nuc |
| ZMIZ2 | 264- GVKRTYSEVYPGQQYL -279 | Nuc |
| TPX2 | 312- GKKRTFDETVSTYVPL -327 | Nuc |
| TPR | 204- KTDELLALGREKGNEI -219 | Cyto |
| SV40 | 47- PKKKRKV -53 | Nuc |
| MYC | 320- PAAKRVKLD -328 | Nuc |
| NEG | DPFKLMDIAACTYKED | Cyto |

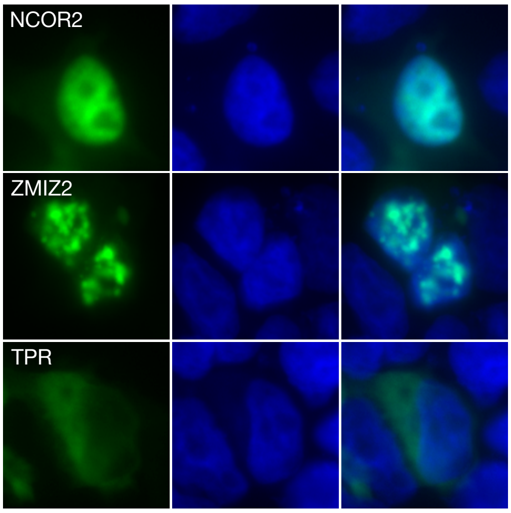

SI Fig. S8

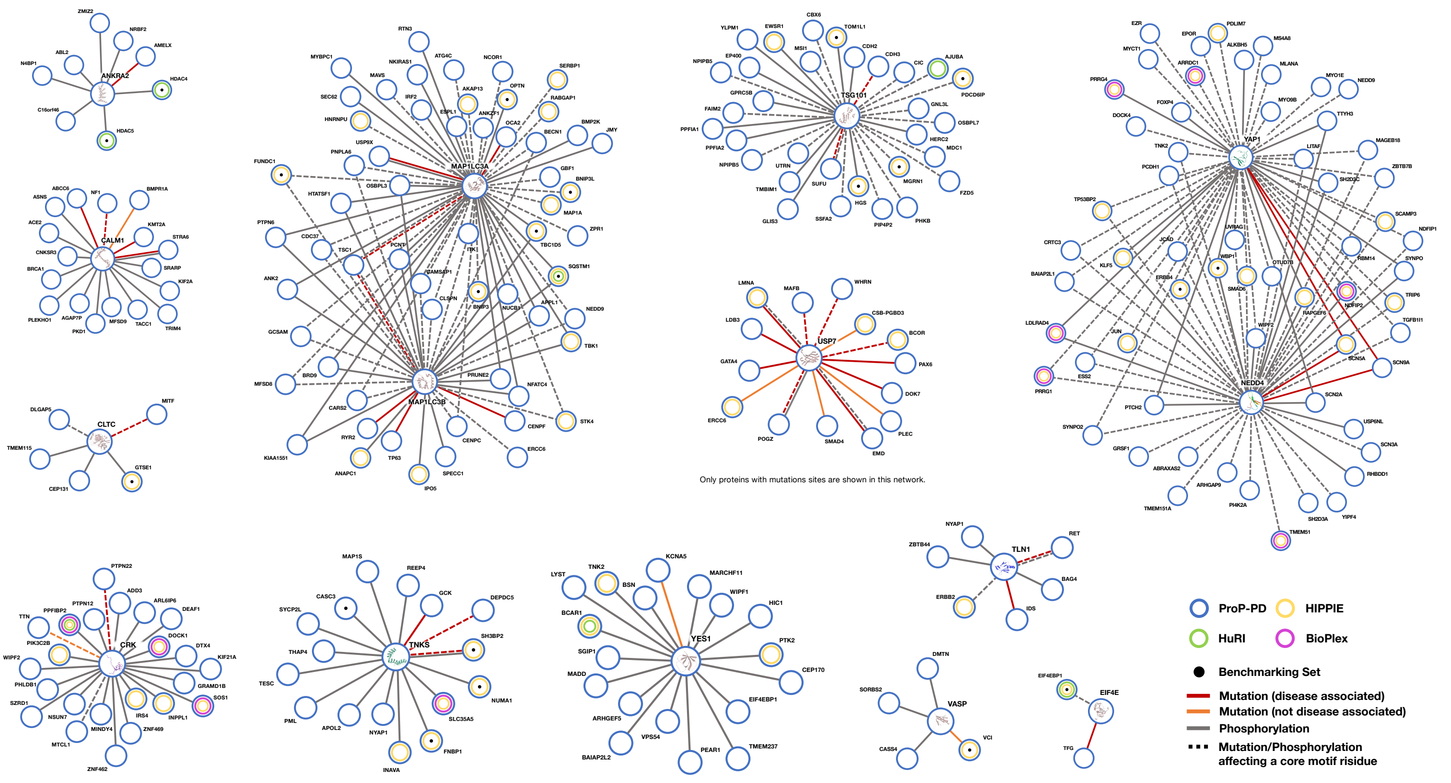


#### ANKRA2

| | |
|---|---|
| AMELX | 169-PMLPDLTLEAWPSTDK-184 |
| HDAC4 | 344-PLYTSPSLPNITLGLP-359 |
| HDAC5 | 364-YTSPSLPNISLGLQAT-379 |
| C16orf46 | 371-VFPRPVLPSLTVSRVI-386 |
| N4BP1 | 824-LPQQPHFPLLPALPSL-839 |
| ABL2 | 561-SSSSVVPYLPRLPILP-576 |
| ZMIZ2 | 390-NQEVKSPFLPDLKPNL-405 |
| NRBF2 | 255-DIPIPNLPPLDFPSPE-270 |

#### CALM1

| | |
|---|---|
| ABCC6 | 208-AAFPSKATFWWVSGLV-223 |
| ASNS | 532-WLSHYWMPKWINATDP-547 |
| ACE2 | 1-MSSSSWLLLSLVAVTA-16 |
| NF1 | 779-DTHAKWEQATKLILNY-794 |
| BMPR1A | 438-WEMARRCITGGIVEEY-453 |
| KMT2A | 3689-AWKSLTDKVQEARSNA-3704 |
| STRA6 | 639-WGLAYTLLHNPTLQVF-654 |
| SRARP | 148-STWGTVKDSLKALSSC-163 |
| MFSD9 | 101-CWSDVVGRRSSLLACI-116 |
| PKD1 | 3418-LSWPDLLSDPSIVGSN-3433 |
| AGAP7P | 320-KWPSLATLACTPISSS-335 |
| KIF2A | 603-VSPQLFTFHEAVSQMV-618 |
| BRCA1 | 372-WITLNSSIQKVNEWFS-387 |
| CNKSR3 | 538-TEPSLLVSWFTRLKLL-553 |
| PLEKHO1 | 366-WSQAKRVLQEVRELRD-381 |
| TACC1 | 4-SPWQILSPVQWAKWTW-19 |
| TRIM4 | 334-SASSWPVFSSAWNYFA-349 |

#### CLTC

| | |
|---|---|
| CEP131 | 92-PTEPTDFLMLFEGSPS-107 |
| DLGAP5 | 831-FGGNLITFSPLQPGEF-846 |
| MITF | 252-ANTLPVSGNLIDLYGN-267 |
| GTSE1 | 648-LPLIDFCDTPEAHVAV-663 |
| TMEM115 | 336-APESSLITFEAAPPTL-351 |

#### CRK

| | |
|---|---|
| ADD3 | 12-TPPPPSMPHKERYFDR-27 |
| ARL6IP6 | 80-SPVLPDKRNGIFPAAA-95 |
| DEAF1 | 425-ALAVPPPTPTKAAPPA-440 |
| DOCK1 | 1794-DVADVPPPLPLKGSVA-1809 |
| DTX4 | 311-SGVPTVPVKNLNGSSP-326 |
| GRAMD1B | 269-TKPDASPQLPKKSITN-284 |
| INPPL1 | 139-PPLPPRSGSTSISAPT-154 |
| IRS4 | 823-NDNSEYVPMLPGKFLG-838 |
| KIF21A | 1209-KELSPPPGLPSKIGSI-1224 |
| MINDY4 | 286-TASSPPHLPSKRLPPW-301 |
| MTCL1 | 1395-LMDISPFLPEKGLPST-1410 |
| NSUN7 | 580-LSETVTKPPLPQKNTA-595 |
| PHLDB1 | 943-PHSSPPPLPAKASRQL-958 |
| PIK3C2B | 152-KKLSPPPLPPRASIWD-167 |
| PPFIBP2 | 415-PFLAEHKYPTLPGKLS-430 |
| PTPN12 | 672-DSPPPLPERTPESFVL-687 |
| PTPN22 | 612-EIPPPLPVRTPESFIV-627 |
| SOS1 | 1210-SPPLLPPREPVRTPDV-1225 |
| SZRD1 | 76-NSTSRPTLPVKSLAQR-91 |
| TTN | 10960-PEVPKKPVPEKKVPVP-10975 |
| WIPF2 | 268-PTNESAPELPQRHNSL-283 |
| ZNF462 | 283-TNLPDVPNKSAPSPTS-298 |
| ZNF469 | 1511-APSLPGKGSGCSVALM-1526 |

#### EIF4E

| | |
|---|---|
| TFG | 97-LESSQVKYLRRELIEL-112 |
| EIF4EBP1 | 52-IIYDRKFLMECRNSPV-67 |

#### MAP1LC3A/B

| | |
|---|---|
| AKAP13 | 597-ISDGLEPYTLLAAGIG-612 |
| ANKZF1 | 526-LLSAPLGSGGFTLLHA-541 |
| BECN1 | 92-ESANSFTLIGEASDGG-107 |
| BMP2K | 1126-VPFTELVVQSITPHQS-1141 |
| BNIP3L | 32-LNSSWVELPMNSSNGN-47 |
| ESPL1 | 1519-TPAPGPEAASGEWELL-1534 |
| GBF1 | 1334-RSATDADVVNSGWLVV-1349 |
| MAP1A | 1784-EEDKLTRSPFEIISPP-1799 |
| USP9X | 1731-YECEESFTTLNVDIRN-1746 |
| OCA2 | 665-ADIHDFEIILHRVEWA-680 |
| ZPR1 | 189-ASPFTLIIDDPSGNSF-204 |
| HNRNPU | 466-YFPIPEEYTFIQNVPL-481 |
| NKIRAS1 | 158-IEPFTLLASKLSQPQS-173 |
| SEC62 | 359-NGNDFEMITKEELEQQ-374 |
| TP63 | 51-SPEVFQHIWDFLEQPI-66 |
| IRF2 | 148-SNGVSDLSPEYAVLTS-163 |
| MAVS | 430-SPFSGCFEDLAISAST-445 |
| OPTN | 168-LNSSGSSEDSFVEIRM-183 |
| RABGAP1 | 16-SVSTLNSEDFVLVSRQ-31 |
| JMY | 942-VLREGFTLLPDTDPLT-957 |
| MYBPC1 | 28-AGTTPAKDWTLVETPP-43 |
| SERBP1 | 16-RFDQLFDDESDPFEVL-31 |
| NCOR1 | 2423-EPAPLLSAQYETLSDS-2438 |
| ATG4C | 443-EKKQLKRFSTEEFVLL-458 |
| RTN3 | 236-DDVIEKDSPESPFEVI-251 |

##### MAP1LC3A

| | |
|---|---|
| FUNDC1 | 8-PQDYESDDDSYEVLDL-23 |
| BRD9 | 577-DVTHDPYEFLQSPEPA-592 |
| MFSD8 | 16-DTPGSREWDILETEEH-31 |
| NFATC4 | 242-GPEDSWLLLSAPGPTP-257 |
| STK4 | 428-PQDGDYEFLKSWTVED-443 |
| CARS2 | 545-STTSTWELLDQRTKDQ-560 |
| TBK1 | 163-ELEDDEQFVSLYGTEE-178 |
| KIAA1551 | 1706-SFSADGFEMLQNPVKD-1721 |
| CENPC | 528-SDWWVVKSEESPVYSN-543 |
| CDC37 | 1-MVDYSVWDHIEVSDDE-16 |
| ANK2 | 1588-VEEEWVIVSDEEIEEA-1603 |
| SQSTM1 | 334-GDDDWTHLSSKEVDPS-349 |
| HTATSF1 | 120-AESGWFHVEEDRNTNV-135 |
| BNIP3 | 80-QGSWVELHFSNNGNGG-95 |
| ITK | 195-NEEYCLLDSSEIHWWR-210 |
| CAMSAP1 | 725-SHDSEPWTLLRQDSDS-740 |
| PCNT | 644-SQELPWVHLQGVQDGD-659 |
| TSC1 | 400-LCHSDDYVHISLPQAT-415 |
| TBC1D5 | 778-DDDSSKDSGFTIVSPL-793 |
| APPL1 | 676-PNQASSEGQFVVLSSS-691 |
| PNPLA6 | 1277-FPSSGFTDLAEIVSRI-1292 |
| PTPN6 | 245-WEEFESLQKQEVKNLH-260 |
| NEDD9 | 619-DGSERSWMDDYDYVHL-634 |
| NUCB1 | 328-DTGEGWETVEMHPAYT-343 |
| CLSPN | 764-SLLTKESSHNSSFELI-779 |
| PRUNE2 | 2722-EWEMLSPQPVQKNMIP-2737 |
| GCSAM | 141-ARSPEDEYELLMPHRI-156 |
| OSBPL3 | 32-QDSWEVVEGLRGEMNY-47 |

##### MAP1LC3B

| | |
|---|---|
| CENPF | 1382-APLDESNSYEHLTLSD-1397 |
| RYR2 | 1340-DSDFEVLMKTAHGHLV-1355 |
| SPECC1 | 325-SPDASDFEHITAETPS-340 |
| ANAPC1 | 702-GSDDDWEYLLNSDYHQ-717 |
| ERCC6 | 1278-DYVLVEAEANRVAQDA-1293 |
| IPO5 | 649-ENMSDDDGWEFVNLGD-664 |

#### TLN1

| | |
|---|---|
| RET | 1051-ALPSTWIENKLYGMSD-1066 |
| ERBB2 | 1238-KGTPTAENPEYLGLDV-1253 |
| IDS | 144-NHTDDSPYSWSFPPYH-159 |
| NYAP1 | 539-DPTVGPLTPLWTYPAT-554 |
| ZBTB44 | 222-VDSSLAFPWTFPFGID-237 |
| BAG4 | 181-TPVSRWIYPQQDCQTE-196 |

#### TNKS

| | |
|---|---|
| APOL2 | 246-IGRISAEGGEQVERVV-261 |
| CASC3 | 144-GERQSGDGQESTEPVE-159 |
| DEPDC5 | 671-SNSRQPGDGMSFLNFS-686 |
| FNBP1 | 513-QDRESPDGSYTEEQSQ-528 |
| GCK | 58-LPTYVRSTPEGSEVGD-73 |
| INAVA | 640-LRRSPDGAPVQVFVPE-655 |
| MAP1S | 642-ESHRSPAEGSERLSLS-657 |
| NUMA1 | 1742-PRTQPDGTSVPGEPAS-1757 |
| NYAP1 | 348-RTPGDGVSRLPVLCHS-363 |
| PML | 524-GPPSPRSPVIGSEVFL-539 |
| REEP4 | 170-PLYLEDQVSHRRPPIG-185 |
| SH3BP2 | 408-PQLPHLQRSPPDGQSF-423 |
| SLC35A5 | 391-RDLSGNLWERSSGDGE-406 |
| SYCP2L | 679-TEERELPEGISTSSLE-694 |
| TESC | 155-RSIADGAMMEAASVCM-170 |
| THAP4 | 163-SQEQAQQALERTPGDG-178 |

#### TSG101

| | |
|---|---|
| CBX6 | 295-LLPETVPSAPSWREP-310 |
| CDH2 | 842-AADNPTAPPYDSLLV-857 |
| CDH3 | 765-ANTDPTAPPYDTLLVF-780 |
| CIC | 892-VPSAPPPKAQSVSPVQ-907 |
| FZD5 | 1-MARPDPSAPPSLLLLL-16 |
| GNL3L | 40-VPSAPHFVHPNDHANR-55 |
| GPRC5B | 384-VLNGGTIPTAPPSHTG-399 |
| HERC2 | 3481-PSAVTPSAPSASARPF-3496 |
| MDC1 | 1156-KTPEPVVPTAPELQPS-1171 |
| MGRN1 | 401-VSPAIPSAPLYEEITY-416 |
| MSI1 | 249-ERTPLPSAPVLPELTA-264 |
| PIP4P2 | 12-LLSASHSGNVTPTAPP-27 |
| SSFA2 | 1151-ASVALTPTAPSRTGSV-1166 |
| SUFU | 11-GPTAPPAPGPTAPPAF-26 |
| TMBIM1 | 1-MSNPSAPPPYEDRNPL-16 |
| UTRN | 867-SQPSAPDFVQRGFDSF-882 |
| YLPM1 | 1523-GSIVRPSAPPARSSVP-1538 |
| AJUBA | 109-FRLEPTAPALSPRSSF-124 |
| EP400 | 46-ASPSAPQSPSYQIQQL-61 |
| FAIM2 | 32-PSAPPSYEEATSGEGM-47 |
| HGS | 344-KSPTPSAPVPLTEPAA-359 |
| NPIPB5 | 556-QLTPLPPSAPPSADDN-571 |
| OSBPL7 | 225-PSAPVIPTHQASVTTE-240 |
| TOM1L1 | 309-EPSAPSQDLLDLSPSP-324 |
| GLIS3 | 610-ERFAPSAPSPHHISPR-625 |
| PHKB | 699-QSSTPSAPELGQQPDV-714 |
| PDCD6IP | 707-KDLQQSIAREPSAPSI-722 |
| EWSR1 | 76-GYTTPTAPQAYSQPVQ-91 |
| PPFIA1 | 942-MSLTSPSAPPTSRTTL-957 |
| PPFIA2 | 956-LAIQEMVSLTSPSAPP-971 |

#### USP7

| | |
|---|---|
| BCOR | 1422-LQQLLPASQSTQLPCS-1437 |
| CSB-PGBD3 | 216-DAEPGPSSLGSMLMPV-231 |
| DOK7 | 351-SLSSYAGSSLDVWRAT-366 |
| EMD | 52-SSSAASSYSFSDLNST-67 |
| ERCC6 | 216-DAEPGPSSLGSMLMPV-231 |
| GATA4 | 349-NSSNATTSSSEEMRPI-364 |
| LDB3 | 342-HTAIASASTTAPASSP-357 |
| LMNA | 607-VGGPISSGSSASSVTV-622 |
| MAFB | 56-SSTPLSTPCSSVPSSP-71 |
| PAX6 | 279-SNTPSHIPISSSFSTS-294 |
| PLEC | 980-SSSGSEAAVPSVCFLV-995 |
| POGZ | 252-TPSTSTTPTATQPTSL-267 |
| SMAD4 | 212-ANFPNIPVASTSQPAS-227 |
| WHRN | 608-QPPSSMPSCSGTVFSA-623 |

#### VASP

| | |
|---|---|
| VCL | 841-DFPPPPPDLEQLRLTD-856 |
| DMTN | 184-PAKIETDYWPCPPSLA-199 |
| SORBS2 | 1005-EDYPDPPIPHSYSSDR-1020 |
| CASS4 | 127-TAQVYEFPDPPTSARI-142 |

#### NEDD4/YAP1

| | |
|---|---|
| MS4A8 | 235-PVTSPPSYSSEIQANK-250 |
| MYCT1 | 219-TPPPPAYESIIKAFPD-234 |
| MLANA | 99-NAPPAYEKLSAEQSPP-114 |
| DOCK4 | 1896-ESKTPPPYSVYERTLR-1911 |
| FOXP4 | 620-SSSPPRLSPPQYSHQV-635 |
| ARRDC1 | 400-ILPPEYSSWGYPYEAP-415 |
| MYO9B | 2142-DPPTYCLPPASGQTNG-2157 |
| MYO1E | 955-APPPPGYHQNGVIRNQ-970 |
| ALKBH5 | 300-SVLPPSYASDRLSGNN-315 |
| NEDD9 | 107-QVPPSYQNQGIYQVPT-122 |
| PRRG4 | 193-VALTRKHSVSPPPPYP-208 |
| EPOR | 493-LIPAAEPLPPSYVACS-508 |
| PDLIM7 | 96-APAADPPRYTFAPSVS-111 |
| EZR | 466-VMTAPPPPPPVYEPV-481 |
| FIGN | 392-SSQSSRALTPPSYSTA-407 |
| NUP214 | 492-SATVTGEPPSYSSGSD-507 |

##### YAP1

| | |
|---|---|
| PRRG1 | 193-PPEYEDIVNSNSASAI-208 |
| BAIAP2L1 | 431-SVVIPPPDYLECLSMG-446 |
| CRTC3 | 436-TEAQAQVSPPPPYPAP-451 |
| ERBB4 | 1045-RSEIGHSPPPAYTPMS-1060 |
| ESS2 | 96-REPPPPYVTPATFETP-111 |
| JCAD | 328-LCLSDPGLEPPVYVPP-343 |
| JUN | 160-SASLHSEPPVYANLSN-175 |
| KLF5 | 316-AEMLQNLTPPPSYAAT-331 |
| LDLRAD4 | 249-EGPPPTYSEVMGHHPG-264 |
| NDFIP2 | 172-PVPPPYSVATSLPTYD-187 |
| OTUD7B | 763-ALLPPPYRVADSYSNG-778 |
| RAPGEF6 | 1521-EGPHTHLKPPDYSVAV-1536 |
| RBM14 | 570-NSTPPPYERTRLSPPR-585 |
| SCAMP3 | 40-VYNPFETREPPPAYEP-55 |
| SCN5A | 1971-TSFPPSYDSVTRATSD-1986 |
| SCN9A | 1947-DATSSTTSPPSYDSVT-1962 |
| SH2D3C | 56-DMVTVPKSPPAYARSS-71 |
| SMAD6 | 269-LCGPESPPPPYSRLSP-284 |
| SYNPO | 560-TAPPTYTETLSTAPLA-575 |
| TGFB1I1 | 32-LTPPPSYGHQPQTGSG-47 |
| TP53BP2 | 864-EEYPPYPPPPYPSGEP-879 |
| TRIP6 | 124-EPPPPPAYRTGSLKPN-139 |
| UVRAG | 512-ERLQYKTPPPSYNSAL-527 |
| WBP1 | 128-EDVVHRPGTPPPPYTV-143 |
| WIPF2 | 244-SGQSLAPPPPPYRQPP-259 |
| ZBTB7B | 466-DAPPHYPPPSTAAASP-481 |
| NDFIP1 | 57-DESGFPKPPSYNVATT-72 |
| PTCH2 | 10-LPPSYTPPARTAAPQI-25 |
| SCN2A | 1964-DMTPSTTSPPSYDSVT-1979 |
| LITAF | 52-DGKGMNPPSYYTQPAP-67 |
| MAGEB18 | 255-EYQQVPNSDPPRYEFL-270 |
| PCDH1 | 1025-NPPKYPSKQVGQPFQL-1040 |
| SYNPO2 | 614-ADFPAPPPYSAVTPPP-629 |
| TNK2 | 623-DWDARPLPPPPAYDDV-638 |
| TTYH3 | 487-ENTPLIGRESPPPSYT-502 |

##### NEDD4

| | |
|---|---|
| ABRAXAS2 | 301-ASDPPPPYSDFHPNNQ-316 |
| GRSF1 | 125-YLEDLPPPPEYELAPS-140 |
| ARHGAP9 | 181-PQPLMSEPPVYCNLVD-196 |
| PI4K2A | 12-RAQPPDYTFPSGSGAH-27 |
| SH2D3A | 224-ASERPPTYCELVPRVP-239 |
| RHBDD1 | 289-RNSPPPYGFHLSPEEM-304 |
| SCN3A | 1960-GSSSTTSPPSYDSVTK-1975 |
| TMEM151A | 433-DTEPLESPPCYEDALY-448 |
| TMEM51 | 229-SIEPLTPPPQYDEVQE-244 |
| YIPF4 | 1-MQPPGPPPAYAPTNGD-16 |
| USP6NL | 625-GLAHPPSYSNPPVYHG-640 |

#### YES1

| | |
|---|---|
| KCNA5 | 73-PGVRPLPPLPEELPRP-88 |
| WIPF1 | 388-NGSTSRALPATPQLPS-403 |
| PTK2 | 362-PQKEGERALPSIPKLA-377 |
| BSN | 2893-VKRTLPSPPPEEAHLP-2908 |
| EIF4EBP1 | 68-TKTPPRDLPTIPGVTS-83 |
| PEAR1 | 928-IRDLPSLPGGPRESSY-943 |
| VPS54 | 24-VDPSKHIRPVPSLPDV-39 |
| ARHGEF5 | 828-SAPGSSRIYRPLPPLP-843 |
| MADD | 731-AAVGVSKPLPSVPPSI-746 |
| BCAR1 | 627-DKTSSIQSRPLPSPPK-642 |
| March11 | 46-PRYLPPLPASPETPER-61 |
| HIC1 | 255-PERPLAERELPPRPDS-270 |
| CEP170 | 643-RTLPQLPNEEKSLESH-658 |
| BAIAP2L2 | 284-ASQLEPDRRSLPRTPS-299 |
| TMEM237 | 16-PPRALPPVPSQDDIPL-31 |
| LYST | 2103-SRSLPAFPTSSLLTQS-2118 |
| SGIP1 | 245-TGTPPPLPPKNVPATP-260 |
| DLGAP2 | 611-KTPPPVPPRTTSKPLI-626 |
| TNK2 | 627-RPLPPPPAYDDVAQDE-642 |
